# Under-sampled KBAs, over-sampled roadsides: integrating historical and contemporary records of Malagasy bees

**DOI:** 10.64898/2026.08.23.746540

**Authors:** Damselle Quesada, Nicolas Leclercq, Leon Marshall, Ciaran E.D. Clark, Aina N. Razakamiaramanana, Nicolas J. Vereecken

## Abstract

Madagascar hosts exceptional biodiversity and endemism, yet its native bee fauna remains poorly characterised. We assembled and cleaned the first comprehensive occurrence dataset for Malagasy bees, reducing 10,721 raw records to 4,071 validated occurrences covering 218 georeferenced species of the 224 checklist species across six families (89.3% endemic).

Despite near-complete checklist coverage, the dataset remains critically sparse for Madagascar’s size, with most species documented by only a few records. Sampling was highly uneven across taxa: most genera were underrepresented while a few were disproportionately sampled due to ecological prevalence, detectability, and collector specialisation. Spatial concentration within limited grid cells amplified these biases.

Temporally, effort varied markedly, with historical peaks driven by individual collectors and a post-2010 shift toward Apidae-dominated records. Spatially, 79.9% of 25×25 km grid cells intersecting Madagascar held no bee records, and 77.1% of records fell within 2.5 km of roads, mirroring global sampling patterns. Sampling clustered near major cities, with common species consistently found near roads and rare species spread across wider distance ranges.

Among 231 Key Biodiversity Areas (KBAs), 68.4% were entirely unsampled; sampled KBAs held only 26.7% of all records, and sampling remained uneven even there, leaving substantial undetected diversity across most sites. These results reveal pervasive temporal, spatial, taxonomic, and collector-driven biases, underscoring the need for targeted surveys within KBAs and beyond roadsides to improve coverage of data-deficient species and strengthen conservation assessments.

## Introduction

Sampling biases have become a pervasive challenge in biodiversity research, particularly for insects, where most regions, taxa, historical periods, and ecosystems remain poorly documented. Consequently, true richness patterns, macroecological inference, and conservation prioritization are fundamentally compromised, creating a fragmented and distorted picture of global diversity and extinction risk (Hughes et al., 2021; Rocha-Ortega et al., 2021). Biological records are often clustered in easily accessible areas, predominantly man-made infrastructure (e.g. roads), obscuring ecological signals and systematically excluding less accessible but ecologically important habitats (Dimson & Gillespie, 2023; He et al., 2025; Hughes et al., 2021; Piccolo et al., 2020). As a result, spatial biases are rarely isolated: they interact with temporal, taxonomic, and collector biases that together shape what gets recorded, where, and by whom (Petersen et al., 2021; Ascher et al., 2020).

Bees are among the most widely studied pollinator groups globally, yet they remain deeply affected by sampling biases that interact with species’ functional traits and spatio-temporal dynamics (Fauviau et al., 2022; Graham et al., 2024; Marshall et al., 2024; Orr et al., 2021; Ostwald et al., 2024). With an estimated 24,705–26,164 species worldwide (Dorey et al., 2026), many regions lack even basic knowledge of their bee diversity (Dorey et al., 2026; Marshall et al., 2024; Orr et al., 2021), including national checklists, the very foundations of this knowledge (see Reverté et al. (2023) for the first fully consolidated continental checklist of Europe published recently). Madagascar is one striking example of this phenomenon: despite being a megadiverse island with remarkable levels of endemism (Marshall et al., 2026), vast areas remain undersampled (Antonelli et al., 2022; Goodman, 2022). Although 231 Key Biodiversity Areas (KBAs) have been established to counter mounting threats from deforestation and other anthropogenic pressures (CEPF, 2022; Rogers et al., 2010), it remains unclear whether these conservation priorities adequately capture the diversity of bees.

The last comprehensive record of Malagasy bees dates back a quarter of a century (Pauly et al., 2001), which compiled information on bee distributions, taxonomy, floral associations, and morphological identification for the bees of Madagascar and the surrounding islands. It remains the single historical baseline for the fauna of Malagasy bees: it is data-rich and invaluable, especially given the absence of physical bee repositories in Madagascar and the likelihood that older collections likely being stored abroad, which slows down further identification and keeps local awareness of Malagasy bee diversity low. This dependence highlights the importance of museum and herbarium specimens, whose rich metadata are indispensable for contextualizing newly collected records (Ficetola et al., 2024): historical data anchor contemporary data for capturing current pressures, events, and the status of taxa, so they can be interpreted meaningfully (Navarro et al., 2025; Turvey & McClune, 2025).

In parallel, citizen science has emerged as a compelling tool for the monitoring of bees, offering opportunities to expand data collection while engaging the public (Aceves-Bueno et al., 2015; McKinley et al., 2017; Vereecken et al., 2021). Platforms such as the Global Biodiversity Information Facility (GBIF) and iNaturalist aggregate vast numbers of records from researchers, citizen scientists, and naturalists. However, concerns persist regarding data accuracy, participation bias, and uneven sampling effort (Burgess et al., 2017; Geldmann et al., 2016). Indeed, public interest often gravitates toward charismatic species or easily accessible locations, potentially reinforcing existing knowledge and spatial gaps (Boakes et al., 2010). Further, data cleaning remains a major challenge for such datasets, requiring careful decisions about which records to retain to ensure the highest possible data quality for analyses (Dorey et al., 2023; Rocchini et al., 2023). Similar scrutiny is also necessary for historical datasets, which may contain outdated or incorrect information that must be corrected before use (Boakes et al., 2010).

Both historical and contemporary datasets have been widely used to address ecological questions, species distribution modelling, and biodiversity monitoring (Isaac & Pocock, 2015; Meyer et al., 2016; Troudet et al., 2017), and combining them is a powerful way to resolve variation across time and space. Their joint interpretation nevertheless demands caution, as heterogeneous sampling effort can distort assessments of species’ habitat associations and ecological tolerances (see Ascher et al., 2020 on bumblebees and climate change). Hughes et al. (2021) emphasized that contemporary sources of species occurrence records such as GBIF unevenly sampled and can skew our understanding toward disturbed or suboptimal environments, and that accessibility biases are pronounced worldwide, shaping how we perceive and interpret biodiversity patterns. In Madagascar, long-standing conservation initiatives and the establishment of numerous protected areas have further shaped where sampling has taken place (Eardley et al., 2009; Goodman, 2022; Rakotoarisoa et al., 2024).

Here, we characterize bee occurrence patterns in Madagascar using 10,721 records representing 220 species compiled from historical and contemporary publicly available datasets (hereafter Malagasy bee dataset). We assess temporal and spatial variation in record numbers and species richness, evaluate spatial sampling biases by quantifying the distance of sampling locations to roads and KBAs, and compare these patterns with global bee occurrence data. Finally, we examine how collector dominance shapes temporal and spatial patterns, including potential taxonomic biases and bycatch effects. We hypothesize that Malagasy bee records exhibit uneven temporal and spatial sampling effort, including a strong footprint of human infrastructure, and peaks in occurrence data that reflect the activity of a small number of dominant collectors. By exposing and characterising these biases, our study aims to improve the interpretation and application of biodiversity data for ecological monitoring and conservation planning in Madagascar.

## Materials and Methods

### Dataset compilation and cleaning

We compiled bee occurrence data from two main sources. First, we digitised the historical occurrence data from Pauly et al. (2001), and we then included records of Malagasy bees from published papers (Smith & Schwarz, 2006; Koch, 2010; Pauly, 2015) as well as from publicly available occurrence records, namely from iNaturalist (using research-grade observations only: quality_grade=research&identifications=any&place_id=7783&taxon_id=630955&verifiable=true&spam=false), and from GBIF (https://doi.org/10.15468/dl.naj2gb). Second, we filtered only through the global publicly available occurrence records from GBIF (https://doi.org/10.15468/dl.5ntqzq) to compare road-distance patterns with Madagascar in this study. The global road data was extracted from the Global Roads Inventory Project (GRIP) (Meijer et al., 2018) and we extracted the Madagascar roads to be used for our distance calculations. The shapefiles of Madagascar KBAs were downloaded from World Database of KBAs (BirdLife International, 2025). Further information of the spatial projections and datasets used are detailed in Supplementary Text 1.

Bee occurrence data was aggregated and curated by removing entries lacking temporal data (missing years) or spatial data (unknown coordinates, outside the administrative boundary of Madagascar, clustered near Antananarivo or biodiversity institutions), and correcting or updating taxonomy (missing names, infraspecific records, misspellings, synonyms, and ambiguous species) using the ‘BeeBDC’ (version 1.3.1) (Dorey et al., 2023) and ‘bdc’ (version 1.1.6) (Ribeiro et al., 2022) packages.

We compiled the most updated checklist of Malagasy bees (see Supplementary 1) through an extensive review of taxonomic literature (Pauly et al., 2001; Eardley et al., 2010; Pauly 2015) and digital sources (DiscoverLife, Atlas Hymenoptera) with consultation from bee taxonomists. We retained the following genera from Pauly et al. (2001): *Hoplonomia* Ashmead, *Madagalictus* Pauly 1984, *Pachyhalictus* (*Archihalictus*) Pauly 1984, and *Zonalictus* Michener 1978, in contrary to the recent changes of *Hoplonomia* Ashmead as a subgenus of *Nomia* Latreille 1804, and *Madagalictus* Pauly 1984, *Pachyhalictus* (*Archihalictus*) Pauly 1984, and *Zonalictus* Michener 1978 as subgenera of *Patellapis* Friese 1909. For the genus *Tetraloniella* Ashmead endemic in Madagascar, we transferred it in *Tetralonia* Spinola, 1838 following Frietas et al. (2023) on the updates of Eucerini phylogeny.

The checklist presented here corresponds to the latest state-of-the art, but ongoing research of the Nomiine bees in the Halictidae family is likely to bring changes in the systematics of bees of Madagascar where selected genera and subgenera will be subject to rearrangements (Silas Bossert and Alain Pauly, pers. comm. July 2026). We removed the species from the checklist that are undetected/mislabeled from Madagascar but recorded in the checklist, retaining only those that have records present in Madagascar. We used the final checklist for correcting the names and synonyms by incorporating updated records for new species. After we removed duplicates from scientific name, coordinates, date, and collector (recordedBy).

We compared all species with occurrence records to determine the degree of completeness of the Malagasy bee dataset for each bee family. For each species, we extracted the dates of the first and last observations along with their associated data sources to monitor the status of Malagasy bees. Further, we retained the information of ‘recordedBy’ to identify the collectors’ expertise that influenced their sampling collection of Malagasy bees.

A taxonomic tree was constructed from the checklist using Linnaean hierarchical classification (superfamily, position, family, subfamily, tribe, genus, subgenus, species) via the *as.phylo()* function from the ‘ape’ package (version 5.8) (Paradis & Schliep, 2019). The tree was visualized as a circular dendrogram using the ‘ggtree’ (version 3.10.1) (Yu, 2025) and ‘ggtreeExtra’ (version 1.18.0) (Xu et al., 2021) packages, with family-level clades highlighted and endemic status mapped as an outer tile layer.

To quantify taxonomic sampling bias at the genus level, we calculated a bias ratio for each genus as the ratio of its observed proportion of records in the dataset to its expected proportion of species in the checklist. A bias ratio greater than 1 therefore indicates overrepresentation, while a value below 1 indicates underrepresentation relative to known diversity.

Cumulative species richness and observation counts (here non-duplicate records, log-transformed for visualization) were plotted over time for each data source, decomposed by bee family, to visualize how each source contributed to the overall dataset through time.

### Spatial calculations

We compared road distances between Malagasy and global bee records, and separately quantified the distance of Malagasy bee records to KBAs. Distances from global roads (were we used to measure the distance from the global bee occurrence data), Malagasy roads (bee occurrence data), and KBAs to each unique bee sampling point were computed by identifying the nearest road or KBA and calculating the minimum Euclidean distance using the ‘sf’ package (version 1.1.0) (Pebesma, 2018; Pebesma & Bivand, 2023). To examine potential spatial biases, we plotted separately the observation density (number of records per cell) and species richness (number of species per cell) relative to the minimum distance to roads using a 25 × 25 km grid for visual inspection. Two additional spatial resolutions (10 × 10 km; 50 x 50 km) were also explored.

Raw observation density and species richness were first plotted to highlight the potential dominance of highly sampled cells. To reduce the influence of these dominant cells and better assess spatial sampling biases, observation density and species richness were rescaled using percent-rank transformations of the ‘scales’ package (1.4.0) (Wickham et al., 2011). This approach allowed us to evaluate whether highly sampled cells were disproportionately located near major roads and cities. We also quantified the number of records within KBAs and identified the KBAs lacking sampling records. After we identified the KBAs with records, we calculated the Chao1 estimated species richness using the ‘vegan’ package (2.7.3) (Oksanen et al., 2026) and compared it against observed species richness, which was further decomposed into endemic and non-endemic species. Furthermore, we plotted the relationship between a species frequency (defined here as the number of occurrence records per species) and its average minimum distance to roads per species to explore how detection bias may lead more recorded species to cluster closer to roads.

To identify the geographic range of Malagasy bees from the compiled dataset, we estimated the Area of Occupancy (AOO) and Extent of Occurrence (EOO) following IUCN (International Union for Conservation of Nature) Redlist Assessment Criterion B (IUCN SSC, 2012) and using the “red” package (version 1.6.3) (Cardoso & Branco, 2025). AOO was calculated by overlaying 2 x 2km grid cells over known occurrences, which provide the actual occupied space by species, while EOO was delimited by a minimum convex polygon encompassing all records to cover the total geographic spread of each species.

Visualizations and plots were produced using the ‘ggplot2’ package (v 3.5.2) (Wickham, 2016). All data manipulations were conducted in RStudio (RStudio Team, 2025) using R version 4.4.3 for Windows (R Core Team, 2025).

## Results

### Malagasy bee records

The filtering and cleaning steps of the compiled dataset reduced the initial pool of 10,721 records to 4,071 cleaned occurrence records, representing 218 species. The sources with the highest record loss after filtering were GBIF (loss of 4,643 records, only 13.3% of records retained) and Pauly et al. (2001) (loss of 1,954 records, 61.4% retained) with duplicates and missing coordinates as the main contributors. Twenty-three records in GBIF were flagged as ambiguous or absent from the checklist, including cf. *Bombus eriophorus* Klug, 1807 and cf. *B. sylvicola* Kirby, 1837, neither of which has known occurrences in Sub-Saharan Africa and for which voucher specimens were not available to us or otherwise validated by experts. Several species were excluded as their known ranges are restricted to neighboring islands, mainland Africa or Europe rather than Madagascar: *Amegilla madecassa* (Saussure, 1890) (African mainland), *Ceratina tabescens* Cockerell, 1912 (Seychelles), *Heriades aldabrana* Cockerell, 1912 (Aldabra), *Megachile chrysopogon* Vachal, 1910 (Congo, Kenya, South Africa, Zambia) and *Sphecodes albilabris* (Fabricius, 1793) (Europe and North Africa) (Eardley & Urban, 2010; Pauly et al., 2001). *Gronoceras felinum* (Gerstäcker, 1858) was also removed following its presence being identified as a labelling error in Pauly et al. (2001). In terms of species richness, Pauly et al. (2001) was largely the dominant source containing 211 of the 218 species in the final cleaned Malagasy bee dataset (Table 1), 99 of which were unique to this source.

**Table 1.** Summary of the datasets filtering applied using BeeBDC. Numbers close in parentheses are the proportion (%) of the cleaned dataset from the original dataset after applying the filters. Species richness (**SR**) is the total number of species of the cleaned dataset.

| Selection/alterations applied | Pauly et al. (2001) | Other publications (2006, 2010, 2015) | GBIF | iNaturalist | Malagasy bee dataset |
| --- | --- | --- | --- | --- | --- |
| *Temporal and spatial filtering* |  |  |  |  |  |
| No year | 503 | 0 | 119 | 0 | 622 |
| No coordinates | 0 | 0 | 1,684 | 0 | 1,684 |
| Near Antananarivo | 158 | 0 | 3 | 3 | 164 |
| Near biodiversity institutions | 0 | 0 | 2 | 3 | 5 |
| Outside country boundary | 2 | 0 | 4 | 2 | 8 |
| Duplicates | 1,268 | 18 | 2,147 | 22 | 3,455 |
| *Taxonomic modifications* |  |  |  |  |  |
| No scientific name | 0 | 0 | 11 | 0 | 11 |
| Species level filtering | 21 | 0 | 650 | 5 | 676 |
| Synonyms | 623 | 0 | 99 | 116 | 838 |
| Spelling | 71 | 0 | 1 | 0 | 72 |
| Ambiguous species* | 2 | 0 | 23 | 0 | 25 |
| *Number of records* |  |  |  |  |  |
| Original dataset | 5,055 | 30 | 5,355 | 281 | 10,721 |
| Cleaned dataset | 3,101<br>(61.4 %) | 12<br>(40 %) | 712<br>(13.3 %) | 246<br>(87.5%) | 4,071<br>(38 %) |
| Species richness | 211 | 10 | 111 | 12 | 218 |

\*Bee species that are incorrectly labelled as occurring in Madagascar but absent from the island, due to misidentification, wrong geographic assignment, or erroneous data entry.

Prior to duplicate removal, the most record-rich species remained heavily inflated by redundant records (Supplementary Table 2). The stingless bee species *Liotrigona mahafalya* Brooks & Michener, 1988 had the highest count (702 records), almost entirely from GBIF (682); followed by *Apis mellifera unicolor* Latreille, 1804 (530; GBIF: 374, iNaturalist: 156), *Cellariella kalaharica* (Cockerell, 1936) (338; GBIF: 308), *Xylocopa calens* Lepeletier, 1841 (332; Pauly et al., 2001: 147, iNaturalist: 104), *Halictus* (*Seladonia*) *jucundus* Smith, 1853 (289; Pauly et al., 2001: 209), and *Lasioglossum* (*Ctenonomia*) *emirnense* (Benoist, 1955) (264; Pauly et al., 2001: 240). The most up-to-date checklist of Malagasy bees, made openly available through this study, is provided in Supplementary Table 1 with Linnaean classification, sources, and synonyms. The corresponding phylogenetic tree is provided in Supplementary Figure 1. It encompasses six families (Andrenidae, Apidae, Colletidae, Halictidae, Megachilidae, and Melittidae) for 224 species with 200 endemic species (89.3%). Of these, 200 (89.3%) had at least one occurrence record in the dataset, with the Apidae and the Halictidae exceeding 90% and all remaining families reaching full representation (Supplementary Table 3). The seven bee taxa lacking any occurrence record are all described in Pauly et al. (2001) were excluded during our cleaning phase: four species (*Austronomia rainandriamampandryi* Pauly, 2001, *Hasinamelissa spinipennis* (Brooks & Pauly), and *Nubenomia luridipes* (Benoist, 1964) lacked any associated records, while the three others (*Ceratina lativentris* Friese, 1905, *Melanempis fulva* Brooks & Pauly, 2001, *Pachymelus flavithorax* Benoist, 1962) had coordinates but lack of associated year and month metadata.

Beyond this near-complete species coverage, records were highly unevenly distributed across genera (median bias ratio: 0.59 ± 0.64), and most genera were underrepresented. Overrepresentation was most severe in species-poor genera that nonetheless accumulated disproportionately many occurrence records: *Apis* (1 species, *Apis mellifera unicolor* Latreille, 1804; bias ratio 18.6), *Thyreus* (1 species, *Thyreus quinquefasciatus* (Smith, 1879); 5.6), *Halictus* Latreille, 1804 (2 species; 5.5), and *Xylocopa* Latreille, 1802 (3 species; 5.4). Among species-rich genera, *Lasioglossum* Curtis, 1833 (12 species; 2.1) and *Megachile* Latreille, 1802 (13 species; 1.7) were also overrepresented, though more moderately (Supplementary Figure 2).

### Temporal effect

To characterize how each source contributed to the overall Malagasy bee dataset, we visualized the cumulative number of species and the log-transformed number of observations (Figure 1). Pauly et al. (2001) holds the earliest records, including pre-1990 observations, and also contributes the most species and observations to the compiled dataset, dominated by Apidae, Halictidae, and Megachilidae (Figure 1a, e, Table 1, and Supplementary Figure 3). In GBIF, species and observation accumulation were likewise dominated by Apidae and Halictidae, with Colletidae and Megachilidae appearing mainly from the early 2000s onward; pre-1990 records largely correspond to digitized museum specimens and incidental collections from broader research projects (Figure 1c, g). iNaturalist, as a recent citizen science platform, contributes few species, predominantly Apidae, with sparse records from Halictidae and Megachilidae recorded from the late 1990s to the present (Figure 1d, h). Post-2000 publication-based datasets added comparatively little, their newly described species being largely restricted to Apidae and Megachilidae (Figure 1b, f). Across the historical record, three distinct peaks in both species richness and observation counts stand out, driven primarily by Halictidae and Apidae (Figure 1i).

**Figure 1.**
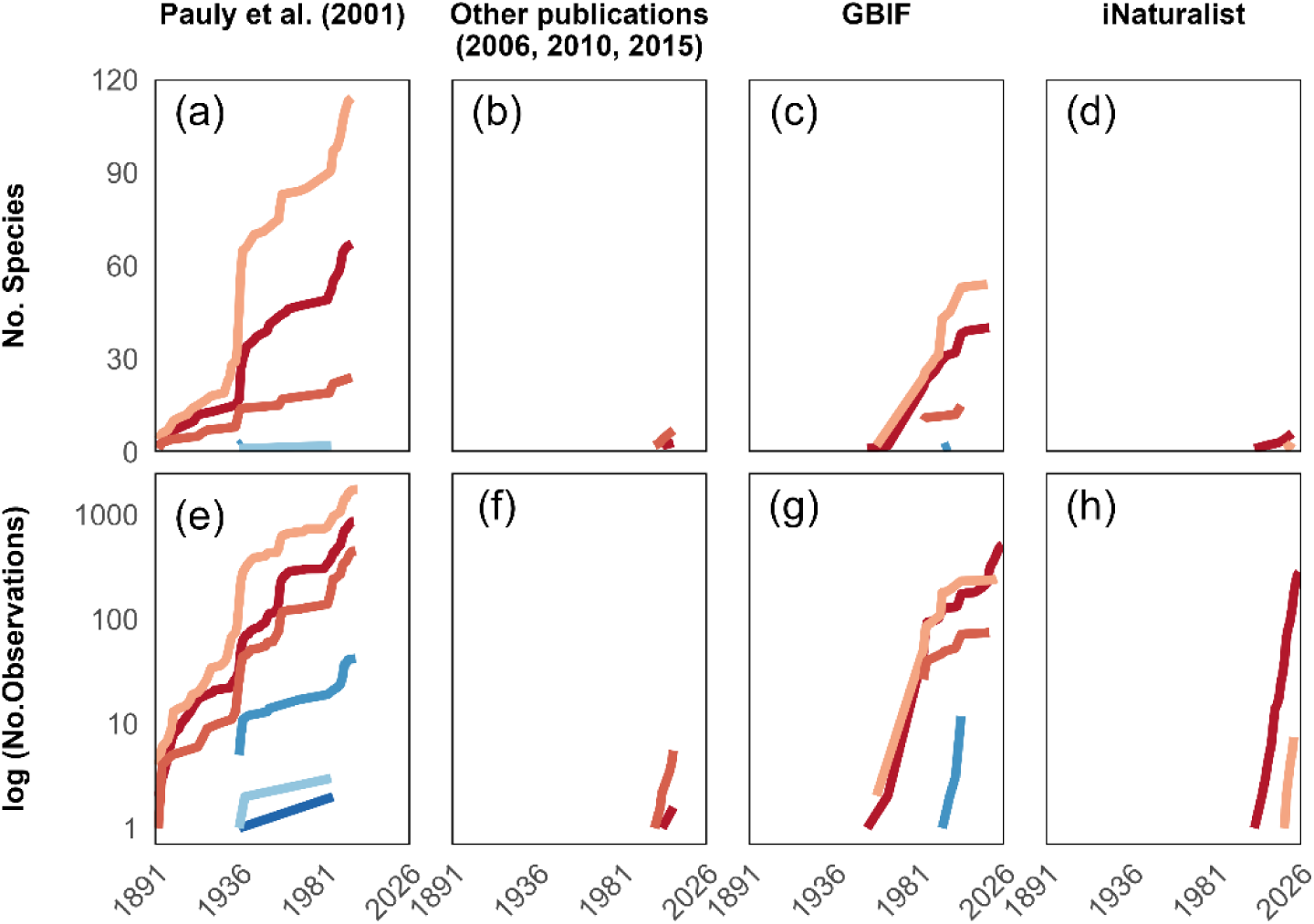
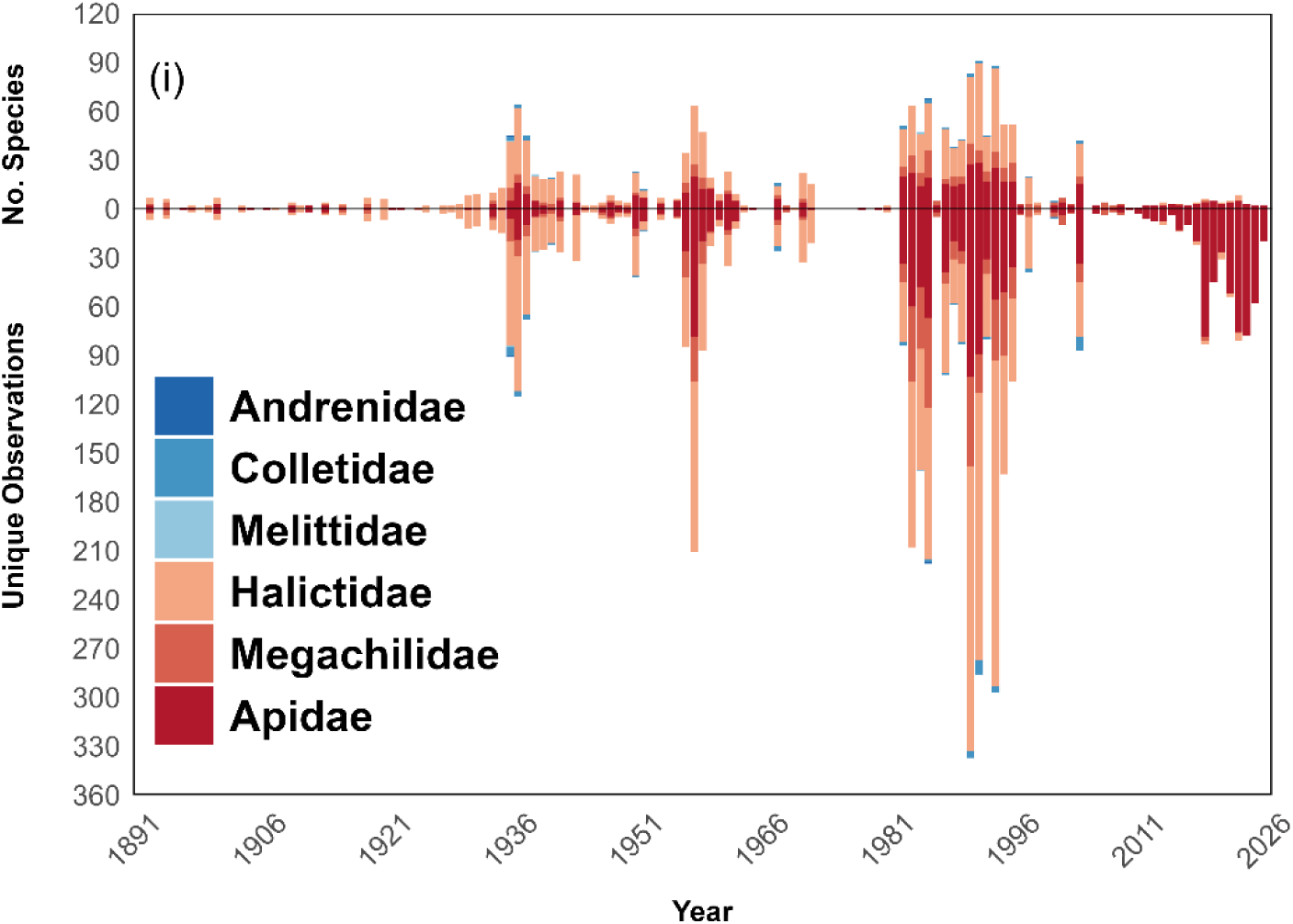
Time-series of Malagasy bee records across six bee families compiled from historical and contemporary datasets. The cumulative number of species from each source (**a-d**), and the log10-transformed number of observations (**e**-**h**). Summary for the Malagasy bee dataset, the upper panel shows the number of species and the lower panel shows the number of observations accounted for by each bee family (**i**).

Over the full compiled dataset, bee records reveal distinct periods of sampling intensity dominated by Apidae, Halictidae, and Megachilidae (Figure 1i; Supplementary Figure 3). Most of these peaks coincide with the activities of specific collectors, many of them entomologists rather than melittologists (i.e., bee scientists), collecting multiple insect taxa at once (Supplementary Table 4) (Pauly et al., 2001). The strong representation of Halictidae in Pauly et al. (2001) reflects Alain Pauly’s specialization in that bee family, whereas GBIF and iNaturalist drove the Apidae and Halictidae peaks recorded between 1981 and 2026. From the early 2010s onward, records have become increasingly dominated by Apidae alone, species that are among the most morphologically recognizable to naturalists and citizen scientists, reinforcing a taxonomic bias toward well-known taxa.

### Spatial effect

Visualizing the bee records in 25 × 25 km grid cells (the 10 × 10 km and 50 x 50 km resolutions are shown in Supplementary Figures 4 & 6) revealed clear patterns of sampling bias. Only 215 of the 1,071 grid cells covering Madagascar contained any records, leaving 79.9% of the island’s surface entirely unsampled. We mapped the raw values of observation density (Supplementary Figure 4e) and species richness (Supplementary Figure 4g) (in red) alongside the minimum distance to roads (in blue). These raw values highlighted a strong imbalance between observation density and species richness across grid cells: with only 2 sampled cells (0.9%) showing high observation counts and 4 cells (1.9%) high species richness, while the vast majority recorded comparatively few observations and species (Supplementary Figure 4e, g). In the percent-rank transformation (Figure 2a), dark purple cells indicate areas with both high observation density and short distances to roads. The near-equal distribution across low, medium, and high percent rank observation density categories is expected by construction; what matters is that across all three density categories, the minimum distance to roads remains consistently low, and that highest-ranked cells are concentrated near major cities and road networks (Figure 2b). We also overlaid the grid with KBAs (Figure 2c), showing that 68.4% of KBAs (158 of the 231) were unsampled.

**Figure 2.**
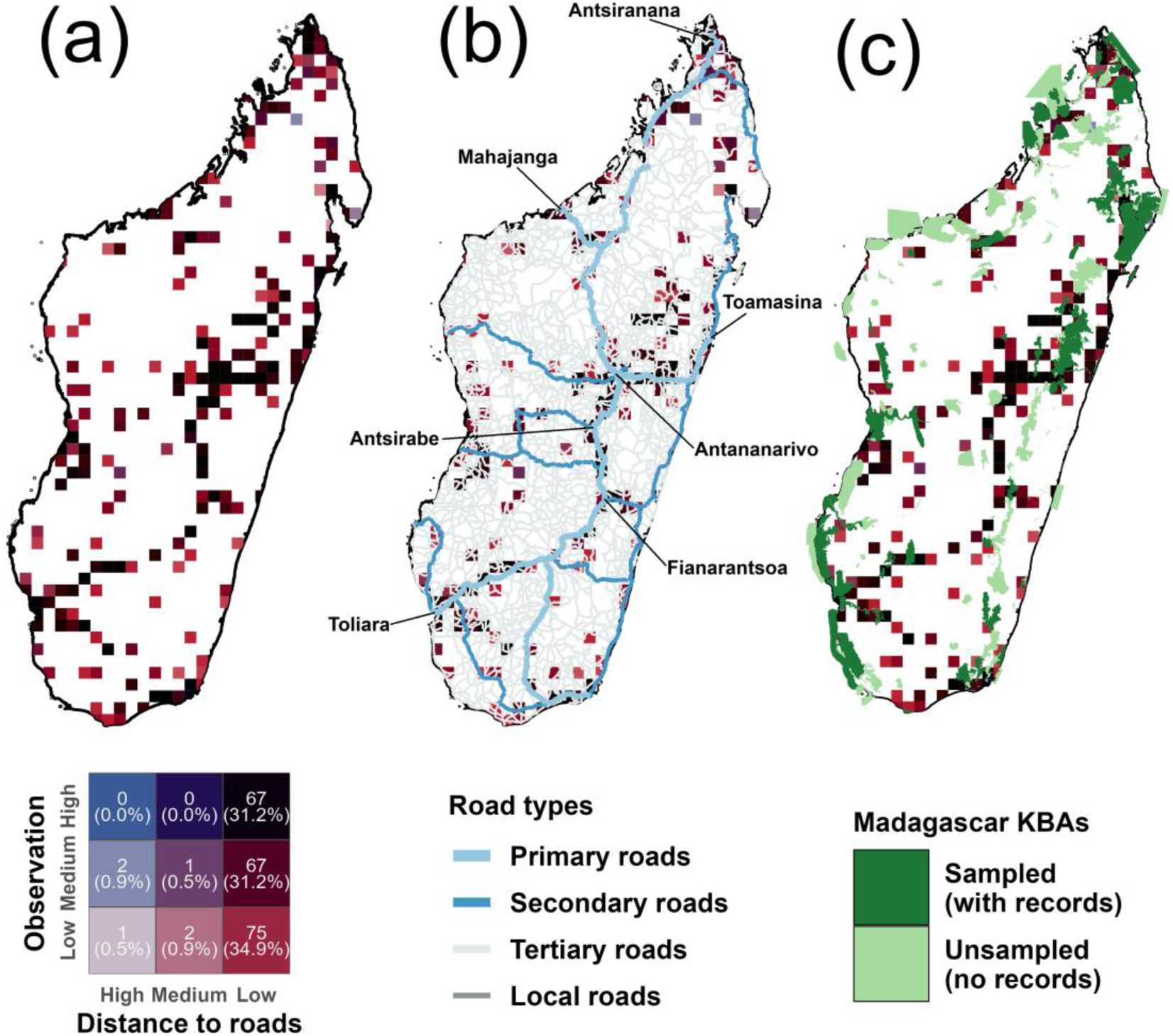
Majority of Malagasy bee records fall near roads and KBAs. (a) Percent rank transformed by observation density in the 25 × 25 km grid. The biplot legend illustrates the mean minimum distance to roads (from high on the left to low on the right, in red) and observation density (from low at the bottom to high at the top, in blue). Cells with both high mean minimum distance to roads and high observation density appear in dark purple. Showing the road networks, major cities (b) and KBAs with (dark green) and without records (light green) (c) falling along the result of biplot bee observation density grid.

Among the 73 sampled KBAs, which collectively encompassed 26.7% of all Madagascar bee records (Supplementary Table 4), sampling effort was highly uneven, with a median of 5 records and 3 species per KBA (mean: 14.9 ± 24.2 records; 7.7 ± 10.6 species). Only 18 KBAs harboured more than 10 species, and the ten most species-rich KBAs accounted for a disproportionate share of total records, led by Ranomafana National Park (86 records, 45 species), Mangoro-Rianila Rivers (120 records, 44 species), and Antainambalana-Andranofotsy River (96 records, 37 species). The remaining 55 sampled KBAs each yielded fewer than 10 species, with many registering only a handful of records. On average, 49.4% (52 KBAs with endemic records out of 73 KBAs with records) of species recorded within KBAs were endemic. The Chao1 richness estimator further showed that observed richness captured only a fraction of estimated true richness across sampled KBAs (median Chao1/observed ratio: 1.21 ± 0.31), with the largest estimation gaps in Sihanaka Forest (30 observed vs. 233 estimated, highly unstable extrapolated richness estimate since 233 more than the total bee fauna of Madagascar), Baie de Diego (16 vs. 69), and PK32-Ranobe (19 vs. 65), sites where the high proportion of singletons suggests that sampling effort was insufficient to capture the full local diversity. Conversely, the 35 KBAs where Chao1 equalled observed richness almost exclusively recorded only one to three species, indicating that sampling was too sparse to reveal any undetected diversity (Supplementary Figure 5).

We compared mean and median distances to roads and KBAs between Malagasy and global bee occurrence records (Figure 3a). Although mean distances to roads were similar between datasets (Madagascar: 1.98km; Global: 1.77km), median distances differed substantially (Madagascar: 1.54km; Global: 0.27km) (Supplementary Table 4). This gap arises because 76.9% of global bee records fall within 1km of roads compared to 40.4% of Malagasy bee records, showing a stronger sampling concentration near roads at the global scale. By contrast, 77.1% of Malagasy records lie within 2.5km of roads, and remain tightly aligned along them. Distances to KBAs were substantially larger, with Malagasy records showing a mean distance of 10.60km and a median of 1.70km (Figure 3a and Supplementary Table 4).

**Figure 3.**
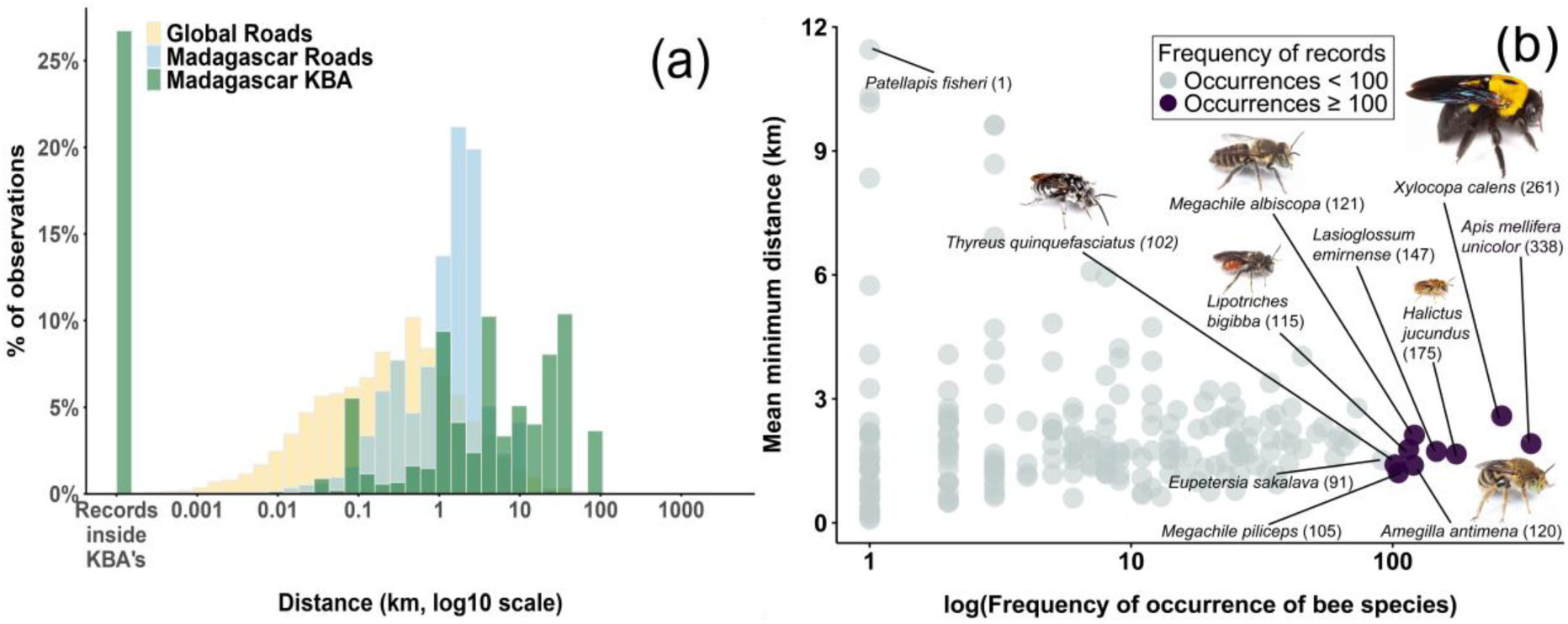
Spatial patterns of Malagasy bees relative to road distance. **(a)** Distribution of Malagasy roads, Malagasy KBAs, and global roads, showing the proportion of records falling below each distance threshold and the corresponding percentage of total observations. **(b)** The spread of the bee species observed across the mean minimum distance (km) to roads. Showing the frequency of occurrence of bee species had been highly observed (purple), equal to or more than 100 occurrences, of which six species had actual and recent field photographs taken by NJ Vereecken available at https://www.flickr.com/photos/nico_bees_wasps/. While the occurrences with less than 100 occurrences (grey).

We also found that rare species occurred across a wide range of distances from roads, whereas most common species were consistently recorded within 3km (Figure 3b). Among the most frequently recorded species (≥100 occurrences), all showed mean distances below 3km, including *A. mellifera* (338 records), *X. calens* (261), *H. jucundus* (175), *L. emirnense* (147), *A. antimena* (121), *M. albiscopa* (120), *L. bigibba* (115), *M. piliceps* (105), and *T. quinquefasciatus* (102). In contrast, the rarest species were recorded at substantially greater distances, with *P. fisheri* representing the most extreme case at up to 12km from the nearest road.

For the 218 species with georeferenced records, we calculated the Area of Occupancy (AOO) and the Extent of Occurrence (EOO) following IUCN guidelines. However, half of all species (52%) returned an EOO below 5,000km², including 103 (41.5%) with a null value, reflecting insufficient spatial coverage to delineate a meaningful range polygon rather than genuinely restricted distributions. AOO remained extremely low throughout (4–504 km²), consistent with the spatially clustered sampling documented above (Supplementary Table 1). These estimates should therefore be interpreted as a first approximation: considerably more spatially distributed records will be needed before robust range assessments can be made for most of Malagasy bee species.

## Discussion

Our study reveals that Malagasy bee occurrence records are strongly clustered near roads, while many KBAs remain poorly sampled or hold no bee records at all. Drawing on a compiled historical and contemporary dataset, we identify several interacting forms of sampling bias, including temporal, spatial, and collector-related biases, alongside erroneous entries in the raw data. Even after cleaning removed more than half of the original records, these biases remained clearly detectable, underlining their potential to distort ecological analyses. Beyond bias, the sheer scarcity of data is in and of itself a major concern: 4,071 cleaned, georeferenced records confined to roughly 20% of Madagascar’s surface represents a critically insufficient baseline for an island of this size and ecological complexity, with many species known from a single record alone.

Compiled datasets inevitably contain errors unless carefully scrutinized and passed through appropriate data cleaning steps, a common challenge when dealing with historical and contemporary insect data where temporal, spatial, and taxonomic biases must be addressed through explicit filtering and modelling approaches (Davis et al., 2023; Dorey et al., 2023). For Pauly et al. (2001), the main taxonomic issue was that many of the names listed have since been synonymized, requiring systematic reconciliation.

Before duplicate removal, several species showed disproportionately high record counts, driven by a combination of colonial biology, morphological conspicuousness, and habitat generalism, traits that increase detectability in roadside-concentrated sampling and bycatch contexts (Michener, 2007). Such imbalances can inflate observation density estimates and distort comparisons across species. Despite the large number of raw GBIF records, the majority were duplicates, and both GBIF and iNaturalist contributions highlight the absence of any dedicated, bee-focused survey in Madagascar. A further credibility concern arose from GBIF records attributing specimens to *Bombus* Latreille, 1802 a genus with no confirmed presence in Madagascar, sub-Saharan Africa, or neighbouring islands (Michener, 2007; Pauly et al., 2001). These misidentifications likely stem from DNA-based comparisons against a *Bombus*-dominated reference library, underscoring that barcoding alone is insufficient for reliable identification (Janko et al., 2024) and that rigorous cleaning is essential to avoid erroneous conclusion, and that field identification and photographs could improve identification accuracy and strengthen species validation (Herrera-Mesías et al., 2022; C. D. Smith et al., 2024).

### Spatio-temporal variation in bee recording and species richness in Madagascar

The temporal variation of the number of observations and species richness highlights how sampling effort has fluctuated unevenly over the past century. Distinct peaks indicate periods of targeted collecting activity separated by years with little to no sampling, a pattern typical of long term occurrence datasets, where quality, quantity, and spatial coverage vary substantially across periods (Díaz-Calafat et al., 2024).The dominance of Apidae, Halictidae, and Megachilidae in these peaks is consistent with their high species richness in Madagascar (Pauly et al., 2001), while the near-absence of Andrenidae, Colletidae, and Melittidae reflects both their genuine rarity on the island and the fact that their global distributions are largely centred outside the Afrotropical region (Almeida et al., 2023; Bossert et al., 2022; Michez et al., 2009). This family-level imbalance is further echoed at the genus level, where sampling was highly uneven and most genera underrepresented. The overrepresentation of specific genera and species primarily reflects their ecological prevalence: all are common and polylectic species that tolerate a wide range of habitats making them the most likely to be encountered by collectors sampling along accessible areas (i.e. typically roads).

Additional factors further impact their detectability: cleptoparasitic relationships structurally inflate counts for both host and parasite pairs (*T. quinquefasciatus – A. antimena*; *E. sakalava – L. emirnense*), large body size makes *X. calens* and *Megachile* spp. conspicuous by-catch targets, and collector specialization (e.g. A. Pauly) further inflates Halictidae records. This mirrors global trends where common, generalist and/or managed species attract disproportionate research attention relative to their diversity, with *A. mellifera unicolor* as the most outstanding example (Nesbit et al., 2026). In contrast, rare species such as *P. fisheri* are seldom encountered, may due to their association with more pristine habitats and smaller body sizes that limit foraging ranges (Greenleaf et al., 2007) which reduce detectability.

The strong clustering of observations around major cities and road networks reflects a broader “curse of accessibility” common to many opportunistic biodiversity records. The pattern is already widely documented elsewhere: collectors and observers disproportionately sample areas that are easy to reach, resulting in spatially biased datasets that overrepresent urban and peri-urban environments (Hughes et al., 2021). For example, Jingu & Ogawa (2025) showed that observation hotspots in Japan’s “public wayside forests” were tightly linked with proximity to publicly maintained roads and high-traffic urban areas. The behavioural origins of this bias are further clarified by the decision-based framework of Arazy & Malkinson (2021), which shows that accessibility effects arise primarily at the monitoring stage (i.e., when observers decide where to go), and are reinforced by variation in expertise and willingness to upload records. Together, these studies highlight that accessibility-driven clustering comprises elements of both spatial bias and behavioral phenomena, rooted in the choices and constraints of observers/collectors. This effect likely obscures any genuinely ecological interactions between species abundance or richness and proximity to human infrastructure, with important implications for interpreting ecological processes in urban systems. Global syntheses of urban trait syndromes, for instance, reveal consistent filtering of behavioural and life-history traits across taxa in datasets disproportionately concentrated in accessible urban and peri-urban areas (Hahs et al., 2023), and evidence for urban ecological traps may itself be inflated because cities are among the best documented habitats, raising the detectability of trap dynamics relative to less accessible rural or natural areas (Zuñiga-Palacios et al., 2021). This complements our study by showing that accessibility not only structures where biodiversity is observed, but also which ecological patterns and processes that can be inferred from unstructured biodiversity data.

Our AOO and EOO calculations further describe the geographic footprint of these species (Supplementary Table 1), and help explain why some are commonly detected while others remain rare unless sampling extends beyond roadsides. By examining the first and last observation years, we can also determine whether species have been recently recorded, and assess whether their known sampling locations still support detectable populations despite gaps of 10–50 years since the last record. Both AOO and EOO are central to range- and status-based protection under the IUCN framework (IUCN SSC 2012). Taken together, these results represent an important step toward building a contemporary and foundational knowledge of Malagasy bees, and in particular toward more efficient sampling that boosts observation counts for the rarer species.

### Accessibility bias: clustered near roads, sparse within KBAs

Sampling bias relative to roads and KBAs show that accessibility strongly shapes where bee data are collected in Madagascar, echoing global patterns. Records heavily clustered near roads, consistent with a well-documented, global accessibility bias in biodiversity data: Hughes et al. (2021) found that 41–65 % of terrestrial biodiversity records worldwide fall within 1 km of a road, with comparable patterns reported for insects excluding bees (Díaz-Calafat et al., 2024; Montgomery et al., 2021). Our analysis of global bee occurrences therefore reproduces this road-distance distribution, aligning to a pervasive global trend. Though the trend does not completely coincide for the analysis of Malagasy bee dataset due to its insufficient records across the island. Analogous roadside bias affects the collection of vertebrates and even herbarium plant records (Kadmon et al., 2004; Reddy & Dávalos, 2003).

With more than two-thirds of the cleaned dataset deriving from Pauly et al., (2001), historical (i.e., largely colonial era) collecting routes have disproportionately influenced the distribution of observations across Madagascar. As these historical records are themselves compiled from earlier researchers, museum accounts, and naturalists who documented bees, they reinforce the same spatial signature. Which regions were visited depend strongly on their accessibility and on the location of the nearest research institutions. In Madagascar, such institutions were few and concentrated in major cities until the early twenty-first century. Limited funding and differing taxonomic priorities further constrained logistics, tending to confine sampling to road accessible areas. A possible factor can also relate to the extent and quality of the road network over time, however was not explored in this study. This constraint is compounded by the sparse and spatially uneven road network of Madagascar, historically concentrated along a few major axes that have changed little over the period spanned by these records; the same accessible corridors were therefore revisited across decades, deepening rather than broadening spatial coverage. Roadside sampling also overrepresents disturbed areas, as the species observed in these locations are more likely to survive in disturbed environments, where such traits are common. In contrast, species that thrive in intact forest or at higher elevations are less likely to be detected, as they have a lower chance of being observed when sampling focuses mainly on roadsides rather than going deeper into forested areas.

KBAs reveal a different but complementary pattern. Of Madagascar’s 231 KBAs, around one-third (73 KBAs) contain at least one bee record, and only about a quarter of all bee observations fall within KBA boundaries (Figure 2C). This means that roughly three-quarters of bee observations were mainly from outside of KBAs. Because roadside sampling favours generalist species (Figure 3B), endemic species that likely occur inside KBAs may simply have gone undocumented (Supplementary Figure 5). This under-representation likely reflects persistent limited entomological activity in KBAs and a historical focus on other taxa (Eardley et al., 2009). Two complementary accessibility biases therefore emerge: (1) a strong pull towards roadsides, consistent with global bee data, and (2) a tendency to sample around rather than within KBAs. Both biases constrain our understanding of bee diversity in Madagascar and reinforce the need for sampling strategies that extend beyond easily accessible areas (Mandeville et al., 2022).

### How are collection biases associated with collector preferences?

Collections by previous researchers were influenced or inspired by Alain Pauly’s work as illustrated from Pauly et al. (2001). The majority of the bee species recorded in his work (Pauly et al., 2001) were from Halictidae as he is a Halictidae expert, but he surveyed all bee species during his stay in Madasgacar deploying traps (Alain Pauly, pers. comm. July 2026). Further, he addressed certain genus such as *Hasinamelissa* Chenoweth & Schwarz, 2008 (previously a group under *Halterapis* Michener 1969) was elusive in traps, similarly to smaller genera like *Braunsapis* Michener 1969 and *Allodape* Lepeletier & Serville 1825, or larger ones such as *Pachymelus* Smith, 1879. This extent further exemplies how we have not fully determined the efficiency of different active vs. passive sampling methods and how they can improve the taxonomic coverage during field surveys especially in unexplored areas of bees like Madagascar, or even in the continental Africa.

The assumption of previous researchers that the areas Alain Pauly sampled were the best places to find bees, resulting in the same localities being revisited again and again. We refer to this pattern as the “*founder-collector effect*”, and involves both aligning new fieldwork with previously productive sites to raise the odds of useful observations, but it also embeds new research into familiarity and preference by encouraging future sampling efforts toward the same places. The Malagasy case shows strongly how a single influential melittologist can shape a region’s records. This phenomenon could be investigated in other regions where influential naturalists have conducted fieldwork, made substantial contributions to the taxonomy and community knowledge of particular insect groups.

Many records carried unknown recordedBy entries (Supplementary Table 4), highlighting the extent of missing collector information in these datasets. We did not filter out records with unknown collectors because the source of each record is still known. Identified collectors were predominantly entomologists, but with very few melittologists among them. The year associated with each record may denote either the collection date or the year of museum accession, depending on the depositor. Overall, the combined dataset used here is dominated by Pauly et al. (2001) (Supplementary Figure 2), a clearly signature the weight of historical collections.

Collector bias also contributes to taxonomic bias in bee collections, particularly within the historical records of Pauly et al. (2001). Although bees are generally considered a charismatic group (Carvalheiro et al., 2013; Gill et al., 2016) and therefore often receive disproportionate research attention (Troudet et al., 2017), this pattern does not hold in Madagascar, where despite their diversity, they remain one of the most understudied insect groups, offering a wide-open, new research frontier. Halictidae was the most frequently observed family in our dataset (Figure 1A, Figure 1E), partly because Alain Pauly collected species mainly within this family, inflating its apparent share. Madagascar is indeed rich in Halictidae, Apidae, and Megachilidae (Supplementary Figure 1; Supplementary Table 3), but more standardised sampling focusing on all three major families might help reevaluate the extent to which Halictidae is genuinely the most species-rich family or simply the most frequently collected due to collector familiarity, potentially leading to the under-representation of other bee families, particularly the Apidae and Megachilidae.

Comparable taxonomic biases are documented in other insect groups, such as in North America, where insect data remain sparse and unevenly distributed (Walsh & Figueroa, 2026). The near-absence of resident melittologists in Madagascar further amplifies this problem, reflecting how bee research has long been overlooked and deprioritized in both research and conservation. Nevertheless, while individual preferences can shape taxonomic bias in participatory platforms, many volunteers are increasingly guided by citizen science initiatives that direct attention toward specific organism groups (Díaz-Calafat et al., 2024). As these efforts expand through the support of researchers and the development of new taxonomic tools, we may begin to observe a broader spectrum of bee species, including those that are currently rarely recorded.

## Conclusion

Using the only compiled record of Malagasy bees currently available, we have shown that its historical and contemporary occurrences are pervaded by interacting temporal, spatial, and collector-driven biases that together translate into taxonomic bias from researcher preferences and the founder-collector effect to citizen scientists’ gravitation toward accessible roadsides. These biases can rarely be eliminated, but they can be made explicit: interpreting such datasets responsibly means keeping their uneven distribution across the landscape firmly in view (see also Ascher et al., 2020).

Several limitations frame these conclusions. Our record-to-road distances are measured against the contemporary road network and therefore approximate, since the network accessible to earlier collectors cannot be reconstructed; we also deliberately set aside environmental covariates, whose role in shaping detectability would be a valuable next step. For KBAs, we did not resolve how far within-KBA records fall from boundaries or how deeply they penetrate the interior: such an analysis would reveal whether observers sample convenient edges or venture inward. Above all, our reliance on pre-existing data makes the case for what these data cannot provide: dedicated, bee-specific fieldwork and long-term monitoring inside KBAs, ideally paired with reference collections, high-quality photographs, new taxonomic tools, and ultimately a DNA barcode library to secure identifications at scale. Recent and ongoing projects like Insect Biome Atlas under local supervision by Madagascar Biodiversity Center based in Antananarivo deployed 50 Malaise traps across 33 sites inside KBAs (Miraldo et al. 2025), and LIFEPLAN project that expands the monitoring from insects to other flora and fauna, providing a new global knowledge of Madagascar’s biodiversity (Hardwick et al. 2024); of which both projects harness country-wide insect surveys and metabarcoding-based research program. Further efforts to fill this gap specifically for bees, has been initiated through the pipeline project BEE-MADA, with focus on bee ecology, education and conservation of Malagasy bees, altogether as practical short-term initiatives to reverse the trends described in this study.

As citizen-science data become a dominant source of rapid biodiversity information (Leachman, 2021; Eardley, 2024) accounting for their embedded biases is essential both to justify biodiversity analyses and to interpret them soundly, minimizing knowledge shortfalls (Marshall et al., 2024). Higher-quality occurrence data will sharpen species distribution models, strengthen ecological inference, and support more accurate conservation planning under climate change (Rahimi & Jung, 2025). For Madagascar’s wild bees, a fauna still largely invisible to conservation (Goodman, 2022), building that evidence base is the necessary first step toward identifying the areas and species most in need of protection.

## Data availability statement

This study used publicly available data as described in the Methods section; no new data were collected.

## Author Contributions

**Damselle Quesada:** Conceptualization; writing – original draft; writing – review and editing; formal analysis; data curation; visualization; resources. **Nicolas Leclercq:** Conceptualization; writing – original draft; writing – review and editing; formal analysis; data curation; visualization; resources. **Leon Marshall:** Conceptualization; writing – review and editing; resources. **Ciaran E.D. Clark:** Conceptualization; writing – review and editing; resources. **Aina N. Razakamiaramanana:** Conceptualization; writing – review and editing; resources. **Nicolas J. Vereecken:** Conceptualization; writing – review and editing; funding acquisition; supervision; resources

## Acknowledgements

We thank Alain Pauly, researcher at Tanzania Wildlife Research Institute (Tanzania) and Royal Belgian Institute of Natural Sciences (Belgium) for the identification and verification of specimens.We are also grateful to all contributors of Malagasy bee datasets in iNaturalist and GBIF, whose efforts made this study possible. We also acknowledge Prof. Jean-François Bastin at the Université de Liège (Belgium) and Prof. Olivia Rakotondrasoa at the Université d’Antananarivo (Madagascar) which led the AGRIFO Project that provided financial support for this study.

## References

Aceves-Bueno, E., Adeleye, A.S., Bradley, D., Tyler Brandt, W., Callery, P., Feraud, M., Garner, K.L., Gentry, R., Huang, Y., McCullough, I., Pearlman, I., Sutherland, S.A., Wilkinson, W., Yang, Y., Zink, T., Anderson, S.E. & Tague, C. (2015) Citizen science as an approach for overcoming insufficient monitoring and inadequate stakeholder buy-in in adaptive management: criteria and evidence. Ecosystems, 18(3), 493–506. Available from: 10.1007/s10021-015-9842-4

Almeida, E.A.B., Bossert, S., Danforth, B.N., Porto, D.S., Freitas, F.V., Davis, C.C., Murray, E.A., Blaimer, B.B., Spasojevic, T., Ströher, P.R., Orr, M.C., Packer, L., Brady, S.G., Kuhlmann, M., Branstetter, M.G. & Pie, M.R. (2023) The evolutionary history of bees in time and space. Current Biology, 33(16), 3409–3422.e6. Available from: 10.1016/j.cub.2023.07.005

Antonelli, A., Smith, R.J., Perrigo, A.L., Crottini, A., Hackel, J., Testo, W., Farooq, H., Torres Jiménez, M.F., Andela, N., Andermann, T., Andriamanohera, A.M., Andriambololonera, S., Bachman, S.P., Bacon, C.D., Baker, W.J., Belluardo, F., Birkinshaw, C., Borrell, J.S., Cable, S., … Ralimanana, H. (2022) Madagascar’s extraordinary biodiversity: evolution, distribution, and use. Science, 378(6623), eabf0869. Available from: 10.1126/science.abf0869

Arazy, O. & Malkinson, D. (2021) A framework of observer-based biases in citizen science biodiversity monitoring: semi-structuring unstructured biodiversity monitoring protocols. Frontiers in Ecology and Evolution, 9, 693602. Available from: 10.3389/fevo.2021.693602

Ascher, J.S., Marshall, L., Meiners, J. & Vereecken, N.J. (2020) Heterogeneity in large-scale databases and the role of climate change as a driver of bumble bee decline. Science (E-Letter). Available at: https://www.science.org/doi/10.1126/science.aax8591 (Accessed 1 July 2026).

BirdLife International (2025) The World Database of Key Biodiversity Areas. KBA Partnership. Available at: www.keybiodiversityareas.org (Accessed 17 November 2025).

Boakes, E.H., McGowan, P.J.K., Fuller, R.A., Chang-qing, D., Clark, N.E., O’Connor, K. & Mace, G.M. (2010) Distorted views of biodiversity: spatial and temporal bias in species occurrence data. PLoS Biology, 8(6), e1000385. Available from: 10.1371/journal.pbio.1000385

Bossert, S., Wood, T.J., Patiny, S., Michez, D., Almeida, E.A.B., Minckley, R.L., Packer, L., Neff, J.L., Copeland, R.S., Straka, J., Pauly, A., Griswold, T., Brady, S.G., Danforth, B.N. & Murray, E.A. (2022) Phylogeny, biogeography and diversification of the mining bee family Andrenidae. Systematic Entomology, 47(2), 283–302. Available from: 10.1111/syen.12530

Bossert, S., Freitas, F.V., Pauly, A., Zhu, G., Crowder, D.W., Orr, M.C., Dorey, J.B. & Murray, E.A. (2025) Phylogeny, antiquity, and niche occupancy of *Trinomia* (Hymenoptera: Halictidae), an Afrotropical endemic genus of Nomiinae. Molecular Phylogenetics and Evolution, 204, 108273. Available from: 10.1016/j.ympev.2024.108273

Burgess, H.K., DeBey, L.B., Froehlich, H.E., Schmidt, N., Theobald, E.J., Ettinger, A.K., HilleRisLambers, J., Tewksbury, J. & Parrish, J.K. (2017) The science of citizen science: exploring barriers to use as a primary research tool. Biological Conservation, 208, 113–120. Available from: 10.1016/j.biocon.2016.05.014

Cardoso, P. & Branco, V.V. (2025) red: IUCN Redlisting Tools (Version 1.6.3). Available from: 10.32614/CRAN.package.red

Carvalheiro, L.G., Kunin, W.E., Keil, P., Aguirre-Gutiérrez, J., Ellis, W.N., Fox, R., Groom, Q., Hennekens, S., Van Landuyt, W., Maes, D., Van De Meutter, F., Michez, D., Rasmont, P., Ode, B., Potts, S.G., Reemer, M., Roberts, S.P.M., Schaminée, J., WallisDeVries, M.F. & Biesmeijer, J.C. (2013) Species richness declines and biotic homogenisation have slowed down for NW-European pollinators and plants. Ecology Letters, 16(7), 870–878. Available from: 10.1111/ele.12121

CEPF (2022) Ecosystem Profile for Madagascar Work Package 1: Identification of important ecosystem services for Ecosystem-based Adaptation (EbA). Available from: https://d29l0tur8ol1gj.cloudfront.net/sites/default/files/kbaplus-analysis-for-madagascar-2022.pdf

Danforth, B.N., Eardley, C., Packer, L., Walker, K., Pauly, A. & Randrianambinintsoa, F.J. (2008) Phylogeny of Halictidae with an emphasis on endemic African Halictinae. Apidologie, 39(1), 86–101. Available from: 10.1051/apido:2008002

Davis, C.L., Guralnick, R.P. & Zipkin, E.F. (2023) Challenges and opportunities for using natural history collections to estimate insect population trends. Journal of Animal Ecology, 92(2), 237–249. Available from: 10.1111/1365-2656.13763

Díaz-Calafat, J., Jaume-Ramis, S., Soacha, K., Álvarez, A. & Piera, J. (2024) Revealing biases in insect observations: a comparative analysis between academic and citizen science data. PLOS ONE, 19(7), e0305757. Available from: 10.1371/journal.pone.0305757

Dimson, M. & Gillespie, T.W. (2023) Who, where, when: observer behaviour influences spatial and temporal patterns of iNaturalist participation. Applied Geography, 153, 102916. Available from: 10.1016/j.apgeog.2023.102916

Dorey, J.B., Fischer, E.E., Chesshire, P.R., Nava-Bolaños, A., O’Reilly, R.L., Bossert, S., Collins, S.M., Lichtenberg, E.M., Tucker, E.M., Smith-Pardo, A., Falcon-Brindis, A., Guevara, D.A., Ribeiro, B., De Pedro, D., Pickering, J., Hung, K-L.J., Parys, K.A., McCabe, L.M., Rogan, M.S., et al. (2023) A globally synthesised and flagged bee occurrence dataset and cleaning workflow. Scientific Data, 10(1), 747. Available from: 10.1038/s41597-023-02626-w

Dorey, J.B., Gilpin, A-M., Johnston, N.P., Esquerré, D., Hughes, A.C., Ascher, J.S. & Orr, M.C. (2026) Estimating global bee species richness and taxonomic gaps. Nature Communications, 17(1), 1762. Available from: 10.1038/s41467-026-69029-4

Eardley, C.D. (2024) Big bees of South Africa: A photographic identification guide to species. SANBI, Pretoria, South Africa.

Eardley, C.D., Gikungu, M. & Schwarz, M.P. (2009) Bee conservation in Sub-Saharan Africa and Madagascar: diversity, status and threats. Apidologie, 40(3), 355–366. Available from: 10.1051/apido/2009016

Eardley, C. & Urban, R. (2010) Catalogue of Afrotropical bees (Hymenoptera: Apoidea: Apiformes). Zootaxa, 2455(1), 1–252. Available from: 10.11646/zootaxa.2455.1.1

Fauviau, A., Baude, M., Bazin, N., Fiordaliso, W., Fisogni, A., Fortel, L., Garrigue, J., Geslin, B., Goulnik, J., Guilbaud, L., Hautekèete, N., Heiniger, C., Kuhlmann, M., Lambert, O., Langlois, D., Le Féon, V., Lopez Vaamonde, C., Maillet, G., Massol, F., et al. (2022) A large-scale dataset reveals taxonomic and functional specificities of wild bee communities in urban habitats of Western Europe. Scientific Reports, 12, 18866. Available from: 10.1038/s41598-022-21512-w

Ficetola, G.F., Manenti, R., Barzaghi, B., Romagnoli, S., Parrino, E.L., Melotto, A., Marta, S., Giachello, S., Balestra, V., Lana, E., Maiorano, L., Pennati, R., Lunghi, E. & Falaschi, M. (2024) Integrating historical and recent data to measure long-term trends of endangered subterranean species. Biological Conservation, 296, 110695. Available from: 10.1016/j.biocon.2024.110695

Freitas, F.V., Branstetter, M.G., Franceschini-Santos, V.H., Dorchin, A., Wright, K.W., López-Uribe, M.M., Griswold, T., Silveira, F.A. & Almeida, E.A.B. (2023) UCE phylogenomics, biogeography, and classification of long-horned bees (Hymenoptera: Apidae: Eucerini), with insights on using specimens with extremely degraded DNA. Insect Systematics and Diversity, 7(4), 3. Available from: 10.1093/isd/ixad012

Geldmann, J., Heilmann-Clausen, J., Holm, T.E., Levinsky, I., Markussen, B., Olsen, K., Rahbek, C. & Tøttrup, A.P. (2016) What determines spatial bias in citizen science? Exploring four recording schemes with different proficiency requirements. Diversity and Distributions, 22(11), 1139–1149. Available from: 10.1111/ddi.12477

Gill, R.J., Baldock, K.C.R., Brown, M.J.F., Cresswell, J.E., Dicks, L.V., Fountain, M.T., Garratt, M.P.D., Gough, L.A., Heard, M.S., Holland, J.M., Ollerton, J., Stone, G.N., Tang, C.Q., Vanbergen, A.J., Vogler, A.P., Woodward, G., Arce, A.N., Boatman, N.D., Brand-Hardy, R., … Potts, S.G. (2016) Protecting an ecosystem service. In: Advances in Ecological Research, Vol. 54, pp. 135–206. Elsevier, Amsterdam. Available from: 10.1016/bs.aecr.2015.10.007

Goodman, S.M. (ed.) (2022) The new natural history of Madagascar. Princeton University Press, Princeton, USA. Available from: 10.1515/9780691229409

Graham, K.K., Glaum, P., Hartert, J., Gibbs, J., Tucker, E., Isaacs, R. & Valdovinos, F.S. (2024) A century of wild bee sampling: historical data and neural network analysis reveal ecological traits associated with species loss. Proceedings of the Royal Society B: Biological Sciences, 291(2028), 20232837. Available from: 10.1098/rspb.2023.2837

Greenleaf, S.S., Williams, N.M., Winfree, R. & Kremen, C. (2007) Bee foraging ranges and their relationship to body size. Oecologia, 153(3), 589–596. Available from: 10.1007/s00442-007-0752-9

Hahs, A.K., Fournier, B., Aronson, M.F.J., Nilon, C.H., Herrera-Montes, A., Salisbury, A.B., Threlfall, C.G., Rega-Brodsky, C.C., Lepczyk, C.A., La Sorte, F.A., MacGregor-Fors, I., Scott MacIvor, J., Jung, K., Piana, M.R., Williams, N.S.G., Knapp, S., Vergnes, A., Acevedo, A.A., Gainsbury, A.M., et al. (2023) Urbanisation generates multiple trait syndromes for terrestrial animal taxa worldwide. Nature Communications, 14, 4751. Available from: 10.1038/s41467-023-39746-1

Hardwick, B., Kerdraon, D., Rogers, H.M.K., Raharinjanahary, D., Rajoelison, E.T., Mononen, T., et al. (2024) LIFEPLAN: A worldwide biodiversity sampling design. PLoS ONE 19(12): e0313353. Available from: 10.1371/journal.pone.0313353

He, J., Yang, W., You, Q., Hu, Q., Cong, M., Tian, C. & Ma, K. (2025) Impacts of road networks on the geography of floristic collections in China. Plant Diversity, 47(3), 403–414. Available from: 10.1016/j.pld.2025.02.001

Herrera-Mesías, F., Bause, C., Ogan, S., Burger, H., Ayasse, M., Weigand, A.M. & Eltz, T. (2022) Double-blind validation of alternative wild bee identification techniques: DNA metabarcoding and in vivo determination in the field. Journal of Hymenoptera Research, 93, 189–214. Available from: 10.3897/jhr.93.86723

Hughes, A.C., Orr, M.C., Ma, K., Costello, M.J., Waller, J., Provoost, P., Yang, Q., Zhu, C. & Qiao, H. (2021) Sampling biases shape our view of the natural world. Ecography, 44(9), 1259–1269. Available from: 10.1111/ecog.05926

Isaac, N.J.B. & Pocock, M.J.O. (2015) Bias and information in biological records. Biological Journal of the Linnean Society, 115(3), 522–531. Available from: 10.1111/bij.12532

IUCN Species Survival Commission (SSC) (2012) IUCN Red List categories and criteria: Version 3.1 (2nd ed.). IUCN, Gland, Switzerland. Available from: https://portals.iucn.org/library/node/10315

Janko, Š., Rok, Š., Blaž, K., Danilo, B., Andrej, G., Denis, K., Klemen, Č. & Matjaž, G. (2024) DNA barcoding insufficiently identifies European wild bees due to undefined species diversity, genus-specific barcoding gaps and database errors. Molecular Ecology Resources, 24(5), e13953. Available from: 10.1111/1755-0998.13953

Jingu, S. & Ogawa, Y. (2025) Effects of accessibility on the records of biodiversity observation by the citizens of urban forests. ISPRS Annals of the Photogrammetry, Remote Sensing and Spatial Information Sciences, X-4/W7-2025, 67–74. Available from: 10.5194/isprs-annals-X-4-W7-2025-67-2025

Kadmon, R., Farber, O. & Danin, A. (2004) Effect of roadside bias on the accuracy of predictive maps produced by bioclimatic models. Ecological Applications, 14(2), 401–413. Available from: 10.1890/02-5364

Koch, H. (2010) Combining morphology and DNA barcoding resolves the taxonomy of Western Malagasy Liotrigona Moure, 1961 (Hymenoptera: Apidae: Meliponini). African Invertebrates, 51(2), 413–421. Available from: 10.5733/afin.051.0210

Marshall, L., Leclercq, N., Carvalheiro, L.G., Dathe, H.H., Jacobi, B., Kuhlmann, M., Potts, S.G., Rasmont, P., Roberts, S.P.M. & Vereecken, N.J. (2024) Understanding and addressing shortfalls in European wild bee data. Biological Conservation, 290, 110455. Available from: 10.1016/j.biocon.2024.110455

Marshall, L., Ascher, J.S., Whittaker, R.J., Orr, M.C., Hughes, A.C., Schrader, J., Weigelt, P., Kreft, H. & Vereecken, N.J. (2026) *Global patterns and drivers of bee diversity and endemism on islands*, preprint, Available from: 10.64898/2026.02.28.708622

McKinley, D.C., Miller-Rushing, A.J., Ballard, H.L., Bonney, R., Brown, H., Cook-Patton, S.C., Evans, D.M., French, R.A., Parrish, J.K., Phillips, T.B., Ryan, S.F., Shanley, L.A., Shirk, J.L., Stepenuck, K.F., Weltzin, J.F., Wiggins, A., Boyle, O.D., Briggs, R.D., Chapin, S.F., et al. (2017) Citizen science can improve conservation science, natural resource management, and environmental protection. Biological Conservation, 208, 15–28. Available from: 10.1016/j.biocon.2016.05.015

Meijer, J.R., Huijbregts, M.A.J., Schotten, K.C.G.J. & Schipper, A.M. (2018) Global patterns of current and future road infrastructure. Environmental Research Letters, 13(6), 064006. Available from: 10.1088/1748-9326/aabd42

Meyer, C., Weigelt, P. & Kreft, H. (2016) Multidimensional biases, gaps and uncertainties in global plant occurrence information. Ecology Letters, 19(8), 992–1006. Available from: 10.1111/ele.12624

Michener, C.D. (2007) The bees of the world (2nd ed.). Johns Hopkins University Press, Baltimore, USA.

Michez, D., Patiny, S. & Danforth, B.N. (2009) Phylogeny of the bee family Melittidae (Hymenoptera: Anthophila) based on combined molecular and morphological data. Systematic Entomology, 34(3), 574–597. Available from: 10.1111/j.1365-3113.2009.00479.x

Miraldo, A., Sundh, J., Iwaszkiewicz-Eggebrecht, E. et al. (2025). Data of the Insect Biome Atlas: a metabarcoding survey of the terrestrial arthropods of Sweden and Madagascar. Sci Data 12, 835. Available from: 10.1038/s41597-025-05151-0

Montgomery, G.A., Belitz, M.W., Guralnick, R.P. & Tingley, M.W. (2021) Standards and best practices for monitoring and benchmarking insects. Frontiers in Ecology and Evolution, 8, 579193. Available from: 10.3389/fevo.2020.579193

Navarro, L.M., Armstrong, C.G., Changeux, T., Frisch, D., Gil-Romera, G., Kaim, D., McClenachan, L., Munteanu, C., Szabó, P., Baranov, V., Blanco-Garrido, F., Camarero, J.J., García, M.B., Grace, M., Izdebski, A., Morueta-Holme, N., Pando, F., Schouten, R., Spitzig, A., et al. (2025) Integrating historical sources for long-term ecological knowledge and biodiversity conservation. Nature Reviews Biodiversity, 1(10), 657–670. Available from: 10.1038/s44358-025-00084-3

Oksanen, J., Simpson, G.L., Blanchet, F.G., Kindt, R., Legendre, P., Minchin, P.R., O’Hara, R.B., Solymos, P., Stevens, M.H.H., Szoecs, E., Wagner, H., Bedward, M., Bolker, B., Borcard, D., Carvalho, G., De Caceres, M., Durand, S., Evangelista, H.B.A., Hannigan, G., et al. (2026) vegan: Community Ecology Package (Version 2.7-5). Available from: 10.32614/CRAN.package.vegan

Orr, M.C., Hughes, A.C., Chesters, D., Pickering, J., Zhu, C-D. & Ascher, J.S. (2021) Global patterns and drivers of bee distribution. Current Biology, 31(3), 451–458.e4. Available from: 10.1016/j.cub.2020.10.053

Ostwald, M.M., Gonzalez, V.H., Chang, C., Vitale, N., Lucia, M. & Seltmann, K.C. (2024) Toward a functional trait approach to bee ecology. Ecology and Evolution, 14(10), e70465. Available from: 10.1002/ece3.70465

Paradis, E. & Schliep, K. (2019) ape: Analyses of phylogenetics and evolution (Version 5.8-1). Available from: https://CRAN.R-project.org/package=ape

Pauly, A. (2015) Nouvelles espèces d’Anthidiini de Madagascar (Hymenoptera: Apoidea: Megachilidae). Belgian Journal of Entomology, 26, 1–30.

Pauly, A., Brooks, R.W., Nilsson, L.A., Apesenko, Y., Eardley, C.D., Terzo, M., Griswold, T., Schwarz, M., Patiny, S., Munzinger, J. & Barbier, Y. (2001) Hymenoptera Apoidea de Madagascar et des iles voisines. Institut Royal des Sciences Naturelles de Belgique, Brussels, Belgium.

Pebesma, E. (2018) Simple features for R: standardized support for spatial vector data. The R Journal, 10(1), 439–446. Available from: 10.32614/RJ-2018-009

Pebesma, E. & Bivand, R. (2023) Spatial Data Science: With Applications in R. Chapman and Hall/CRC, Boca Raton, USA. Available from: 10.1201/9780429459016

Petersen, T.K., Speed, J.D.M., Grøtan, V. & Austrheim, G. (2021) Species data for understanding biodiversity dynamics: the what, where and when of species occurrence data collection. Ecological Solutions and Evidence, 2(1), e12048. Available from: 10.1002/2688-8319.12048

Piccolo, R.L., Warnken, J., Chauvenet, A.L.M. & Castley, J.G. (2020) Location biases in ecological research on Australian terrestrial reptiles. Scientific Reports, 10, 9691. Available from: 10.1038/s41598-020-66719-x

Rahimi, E. & Jung, C. (2025) Investigating the spatial biases and temporal trends in insect pollinator occurrence data on GBIF. Insects, 16(8), 769. Available from: 10.3390/insects16080769

Rakotoarisoa, S., Aly, T.C., Andriajaona, A., Andriamalala, R., Andriamanana, T., Andriamparany, S., Bakarizafy, J.H., Belalahy, R.T., Decampe, A.R., Emeline, H., B., H., Miandrimananana, C., Nasoavina, C., Rabevao, E., Rahajanirina, N., Raharijaona, L., Raherison, M., Rajaonarison, M., Rakotondrabe, A.R. & Ralisata, M. (2024) Madagascar Protected Area Outlook 2024: A conservation assessment of terrestrial Protected Areas in Madagascar. In: Tsiavahananahary, J.T. (ed.), pp. 1–72.

Reddy, S. & Dávalos, L.M. (2003) Geographical sampling bias and its implications for conservation priorities in Africa. Journal of Biogeography, 30(11), 1719–1727. Available from: 10.1046/j.1365-2699.2003.00946.x

Reverté, S., Miličić, M., Ačanski, J., Andrić, A., Aracil, A., Aubert, M., Balzan, M.V., Bartomeus, I., Bogusch, P., Bosch, J., Budrys, E., Cantú-Salazar, L., Castro, S., Cornalba, M., Demeter, I., Devalez, J., Dorchin, A., Dufrêne, E., Đorđević, A., … Vujić, A. (2023) National records of 3000 European bee and hoverfly species: a contribution to pollinator conservation. Insect Conservation and Diversity, 16(6), 758–775. Available from: 10.1111/icad.12680

Ribeiro, B.R., Velazco, S.J.E., Guidoni-Martins, K., Tessarolo, G., Jardim, L., Bachman, S.P. & Loyola, R. (2022) bdc: A toolkit for standardizing, integrating and cleaning biodiversity data. Methods in Ecology and Evolution, 13(7), 1421–1428. Available from: 10.1111/2041-210X.13868

Rocchini, D., Tordoni, E., Marchetto, E., Marcantonio, M., Barbosa, A.M., Bazzichetto, M., Beierkuhnlein, C., Castelnuovo, E., Gatti, R.C., Chiarucci, A., Chieffallo, L., Da Re, D., Di Musciano, M., Foody, G.M., Gabor, L., Garzon-Lopez, C.X., Guisan, A., Hattab, T., Hortal, J., … Malavasi, M. (2023) A quixotic view of spatial bias in modelling the distribution of species and their diversity. NPJ Biodiversity, 2, 10. Available from: 10.1038/s44185-023-00014-6

Rocha-Ortega, M., Rodriguez, P. & Córdoba-Aguilar, A. (2021) Geographical, temporal and taxonomic biases in insect GBIF data on biodiversity and extinction. Ecological Entomology, 46(4), 718–728. Available from: 10.1111/een.13027

Rogers, H.M., Glew, L., Honzák, M. & Hudson, M.D. (2010) Prioritizing key biodiversity areas in Madagascar by including data on human pressure and ecosystem services. Landscape and Urban Planning, 96(1), 48–56. Available from: 10.1016/j.landurbplan.2010.02.002

Smith, C.D., Cornman, R.S., Fike, J.A., Kraus, J.M., Oyler-McCance, S.J., Givens, C.E., Hladik, M.L., Vandever, M.W., Kolpin, D.W. & Smalling, K.L. (2024) Comparing modern identification methods for wild bees: metabarcoding and image-based morphological taxonomic assignment. PLOS ONE, 19(4), e0301474. Available from: 10.1371/journal.pone.0301474

Smith, J.A. & Schwarz, M.P. (2006) Sociality in a Malagasy allodapine bee, *Macrogalea antanosy*, and the impacts of the facultative social parasite, *Macrogalea maizina*. Insectes Sociaux, 53(1), 101–107. Available from: 10.1007/s00040-005-0842-9

Troudet, J., Grandcolas, P., Blin, A., Vignes-Lebbe, R. & Legendre, F. (2017) Taxonomic bias in biodiversity data and societal preferences. Scientific Reports, 7, 9132. Available from: 10.1038/s41598-017-09084-6

Turvey, S.T. & McClune, K. (2025) Expanding the historical baseline: using pre-modern archives to inform conservation from ecological and human perspectives. BioScience, 75(3), 240–250. Available from: 10.1093/biosci/biae127

Vereecken, N.J., Weekers, T., Marshall, L., D’Haeseleer, J., Cuypers, M., Pauly, A., Pasau, B., Leclercq, N., Tshibungu, A., Molenberg, J. & De Greef, S. (2021) Five years of citizen science and standardised field surveys in an informal urban green space reveal a threatened Eden for wild bees in Brussels, Belgium. Insect Conservation and Diversity, 14(6), 868–876. Availbale from: 10.1111/icad.12514

Walsh, W. & Figueroa, L.L. (2026) Data deficiency, taxonomic bias, and economic interests curtail insect and arachnid conservation in the United States. Proceedings of the National Academy of Sciences, 123(10), e2522779123. Available from: 10.1073/pnas.2522779123

Wickham, H. (2016) ggplot2. Springer International Publishing, Cham, Switzerland. Available from: 10.1007/978-3-319-24277-4

Xu, S., Dai, Z., Guo, P., Fu, X., Liu, S., Zhou, L., Tang, W., Feng, T., Chen, M., Zhan, L., Wu, T., Hu, E., Jiang, Y., Bo, X. & Yu, G. (2021) ggtreeExtra: compact visualization of richly annotated phylogenetic data. Molecular Biology and Evolution, 38(9), 4039–4042. Available from: 10.1093/molbev/msab166

Yu, G. (2025) ggtree: Visualization and annotation of phylogenetic trees (Version 3.14.0). Available from: https://bioconductor.org/packages/release/bioc/html/ggtree.html

Zuñiga-Palacios, J., Zuria, I., Castellanos, I., Lara, C. & Sánchez-Rojas, G. (2021) What do we know (and need to know) about the role of urban habitats as ecological traps? Systematic review and meta-analysis. Science of the Total Environment, 780, 146559. Available from: 10.1016/j.scitotenv.2021.146559

