## Supplementary Material for "Under-sampled KBAs, over-sampled roadsides: integrating historical and contemporary records of Malagasy bees"

#### Supplementary Text 1

Spatial data for global roads was divided into seven regions and were reprojected to their respective region-specific projected coordinate systems (in meters) (EPSG:5070 for North America, ESRI:102033 for Central and South America, ESRI:102022 for Africa, EPSG:3035 for Europe, ESRI:102025 for Middle East and Central Asia, ESRI:102028 for South and East Asia, and EPSG:3577 for Oceania) (Meijer et al., 2018). For Madagascar, both boundary and road data were reprojected to the Tananarive (Paris) / Laborde Grid (EPSG:29701).
The global road dataset included five road types: (i) highways, (ii) primary roads, (iii) secondary roads, (iv) tertiary roads, and (v) local roads. Highways were absent from the Madagascar dataset, so only the remaining four road types were used in the calculations for Madagascar (Meijer et al., 2018). Madagascar's administrative boundary shapefiles were obtained from GADM (2025).
Two shapefiles were merged from the WDPA_WDOECM_May2025_Public_MDG_shp_0 and WDPA_WDOECM_May2025_Public_MDG_shp_1 datasets from BirdLife International (2025) to obtain the full set of protected areas overlapping Madagascar's KBAs. Several KBAs include protected area polygons where marine or freshwater components are fused with terrestrial boundaries in the original WDPA geometry. Because these components cannot be reliably separated, there is no clear delimitation in the shapefiles and the aquatic portions are embedded within or contiguous with the terrestrial protected area or national park, thus we retained them rather than removing the marine/freshwater sections.

Seven major cities are based from https://www.geonames.org/mg/largest-cities-in-madagascar.html. The shapefile was extracted from GADM (2025), filtering into administrative boundary 3 (District-level).

**Supplementary Table 1. Madagascar Bee Checklist with the abundance and spatial estimates of AOO and EOO.** The table is provided as a separate file to facilitate data manipulation and reuse.

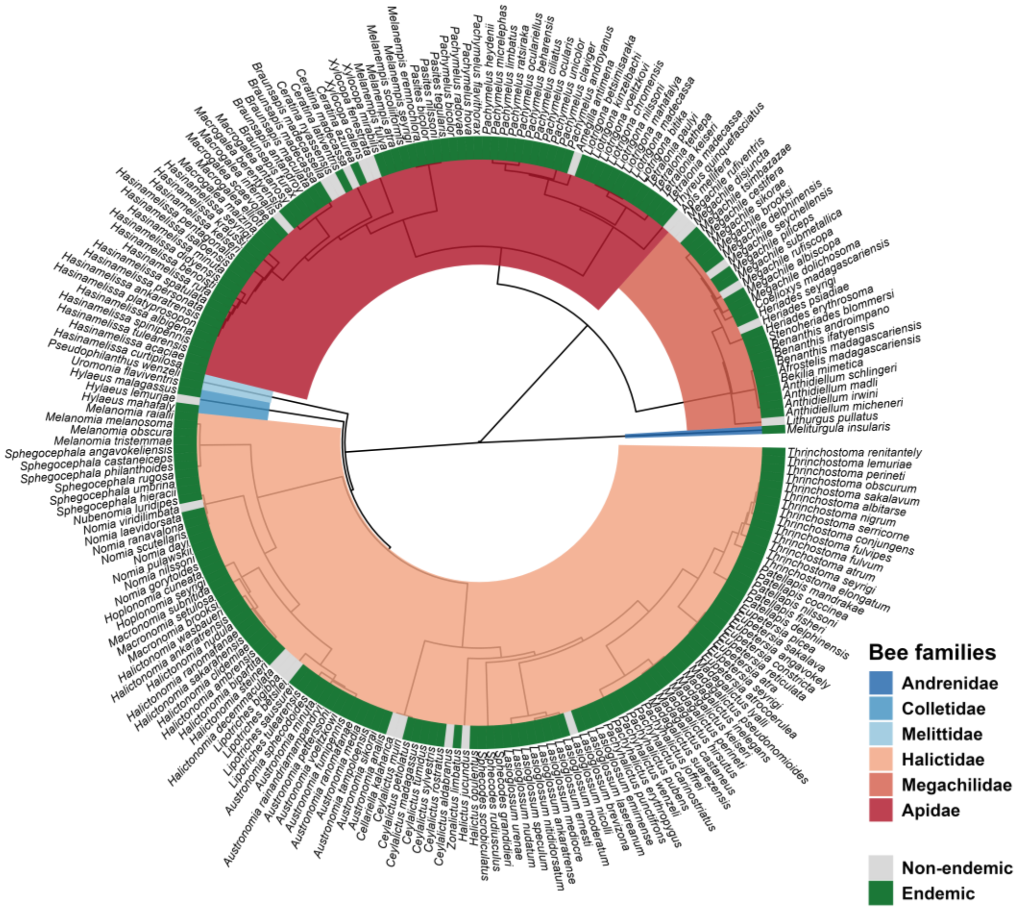


**Supplementary Figure 1. Malagasy bee species circular dendrogram.** The phylogenetic tree of Malagasy bee species is calculated based on Linnaean hierarchical classification (superfamily, position, family, subfamily, tribe, genus, subgenus, species). The bee families are represented by different colors to highlight major lineages. Endemic species are highlighted in green.

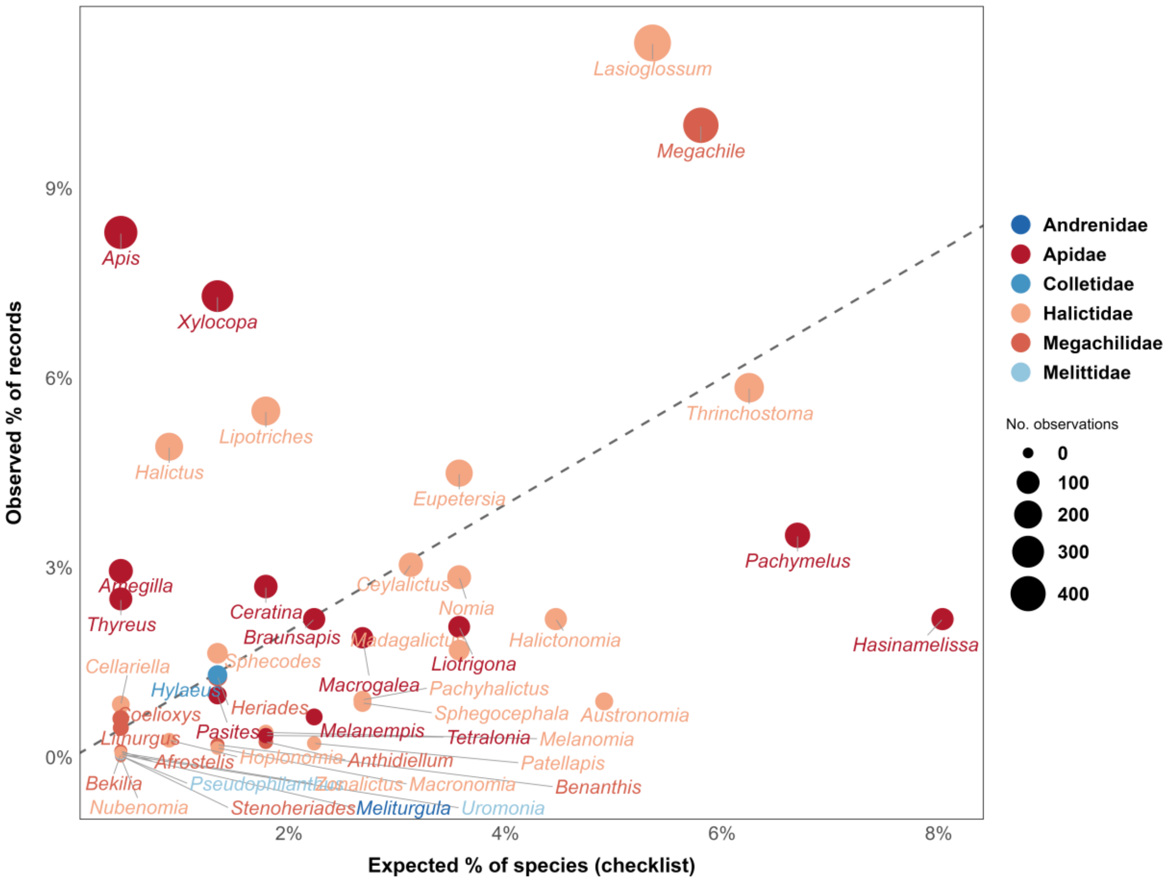


**Supplementary Figure 2. Sampling bias per genus in the Malagasy bee dataset.** Each point represents a genus, plotted against its expected proportion of species from the checklist and its observed proportion of records. The dashed line indicates unbiased sampling (1:1 ratio). Points above the line are overrepresented in the dataset relative to their known diversity, while points below are underrepresented. Point size is proportional to the total number of records. Colours indicate family membership.

**Supplementary Table 2. Checklist completion per bee family**. Checklist completion was calculated for each bee family based on the species currently recorded from Madagascar. The proportion represents how completely the observed species in our datasets match the species listed in the checklist. Completion values therefore reflect the state of available historical and contemporary records. Because Madagascar has no nationwide bee monitoring yet or any bee research-focused project, additional species likely remain undocumented, and the true species richness is expected to be higher than what is captured in the current checklist.

| Bee Families | No. of observed species | No. of species in the checklist | % of completion |
|---|---|---|---|
| Andrenidae | 1 | 1 | 100 |
| Apidae | 70 | 74 | 94.6 |
| Colletidae | 3 | 3 | 100 |
| Halictidae | 114 | 116 | 98.3 |
| Megachilidae | 28 | 28 | 100 |
| Melittidae | 2 | 2 | 100 |
| **Total** | 218 | 224 | 96.9 |

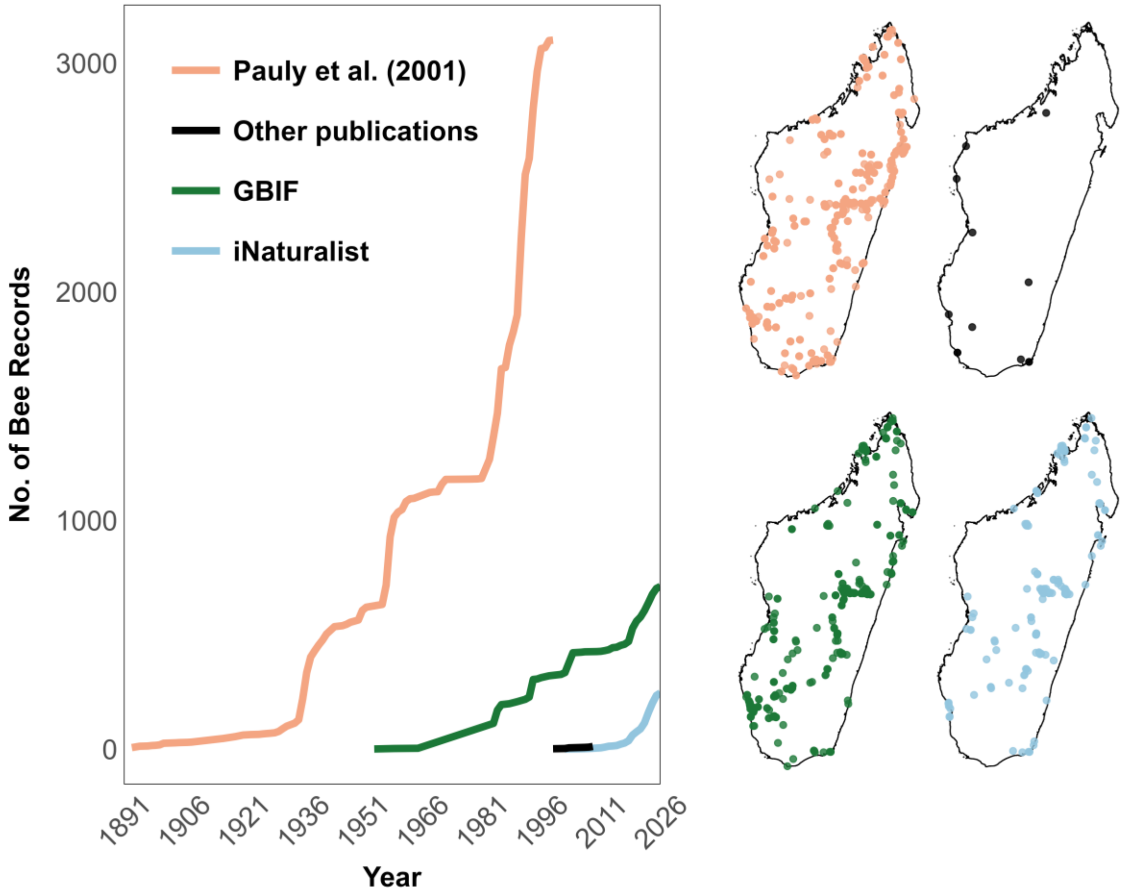


**Supplementary Figure 3. Temporal and spatial distribution of the Malagasy bee dataset.** The right panel shows the cumulative number of bee records across years from the different data sources. Historical records are primarily represented by Pauly et al. (2001), with additional early records contributed by GBIF. The left panel presents the spatial distribution of records from all sources across Madagascar.

**Supplementary Table 3.** The table is provided as a separate file to facilitate data manipulation and reuse. Collectors (RecordedBy) extracted from the Malagasy bee dataset summarized by the years and number of observations (N), following columns are the breakdown of observations of the bee families: Apidae, Halictidae, Megachilidae, Colletidae, Andrenidae, and Melittidae.

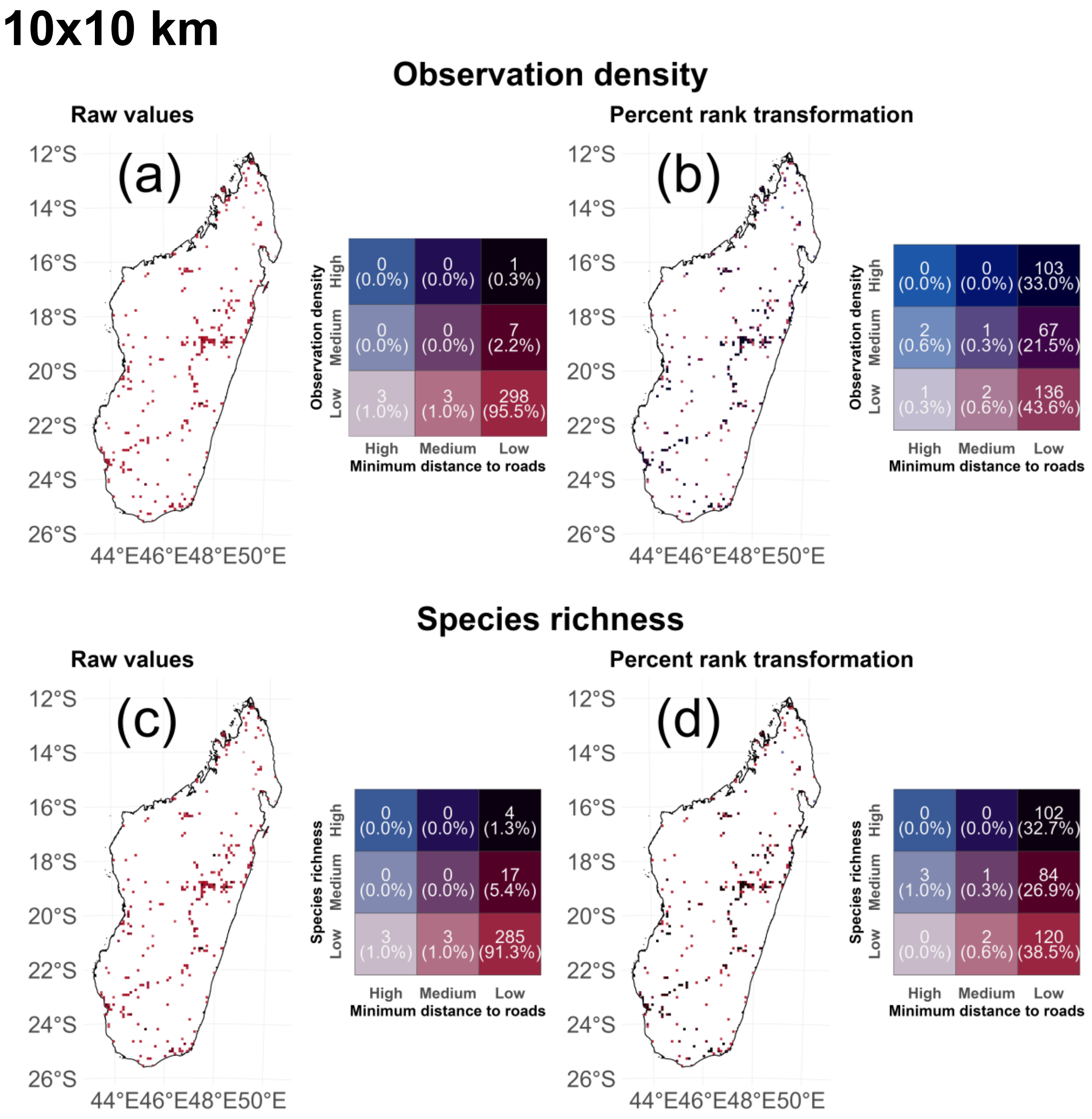

10x10 km
Observation density
Raw values
(a)
12°S
14°S
16°S
18°S
20°S
22°S
24°S
26°S
44°E46°E48°E50°E
Observation density
High
Medium
Low
0 (0.0%)
0 (0.0%)
1 (0.3%)
0 (0.0%)
0 (0.0%)
7 (2.2%)
3 (1.0%)
3 (1.0%)
298 (95.5%)
High
Medium
Low
Minimum distance to roads
Percent rank transformation
(b)
0 (0.0%)
0 (0.0%)
103 (33.0%)
2 (0.6%)
1 (0.3%)
67 (21.5%)
1 (0.3%)
2 (0.6%)
136 (43.6%)
Species richness
Raw values
(c)
0 (0.0%)
0 (0.0%)
4 (1.3%)
0 (0.0%)
0 (0.0%)
17 (5.4%)
3 (1.0%)
3 (1.0%)
285 (91.3%)
Percent rank transformation
(d)
0 (0.0%)
0 (0.0%)
102 (32.7%)
3 (1.0%)
1 (0.3%)
84 (26.9%)
0 (0.0%)
2 (0.6%)
120 (38.5%)

## 25x25 km

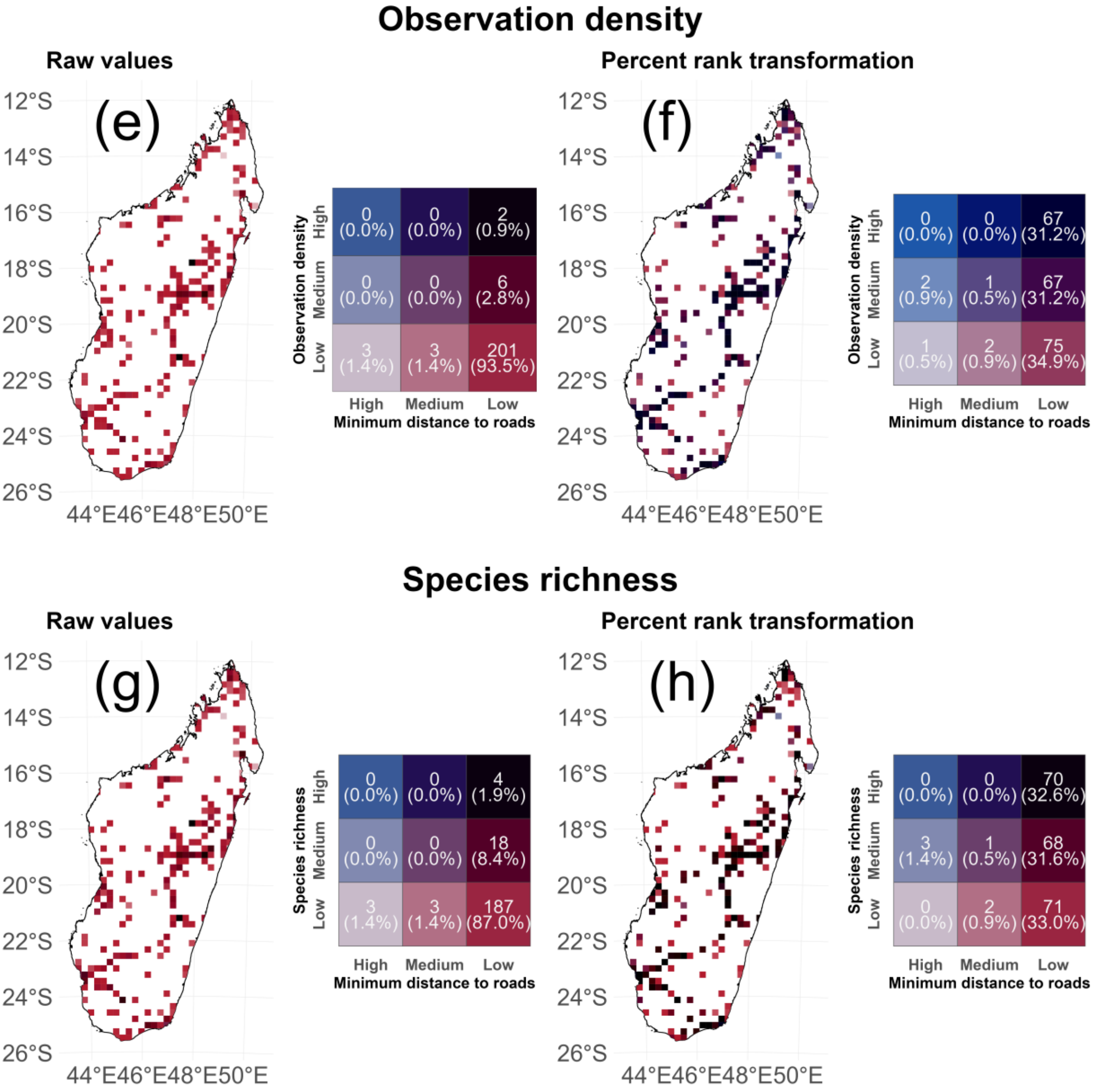

Observation density
Raw values
Percent rank transformation
(e)
(f)
12°S
14°S
16°S
18°S
20°S
22°S
24°S
26°S
44°E46°E48°E50°E
Observation density
High
Medium
Low
0 (0.0%)
0 (0.0%)
2 (0.9%)
0 (0.0%)
0 (0.0%)
6 (2.8%)
3 (1.4%)
3 (1.4%)
201 (93.5%)
High Medium Low
Minimum distance to roads
0 (0.0%)
0 (0.0%)
67 (31.2%)
2 (0.9%)
1 (0.5%)
67 (31.2%)
1 (0.5%)
2 (0.9%)
75 (34.9%)
Species richness
Raw values
Percent rank transformation
(g)
(h)
0 (0.0%)
0 (0.0%)
4 (1.9%)
0 (0.0%)
0 (0.0%)
18 (8.4%)
3 (1.4%)
3 (1.4%)
187 (87.0%)
0 (0.0%)
0 (0.0%)
70 (32.6%)
3 (1.4%)
1 (0.5%)
68 (31.6%)
0 (0.0%)
2 (0.9%)
71 (33.0%)

# 50x50 km

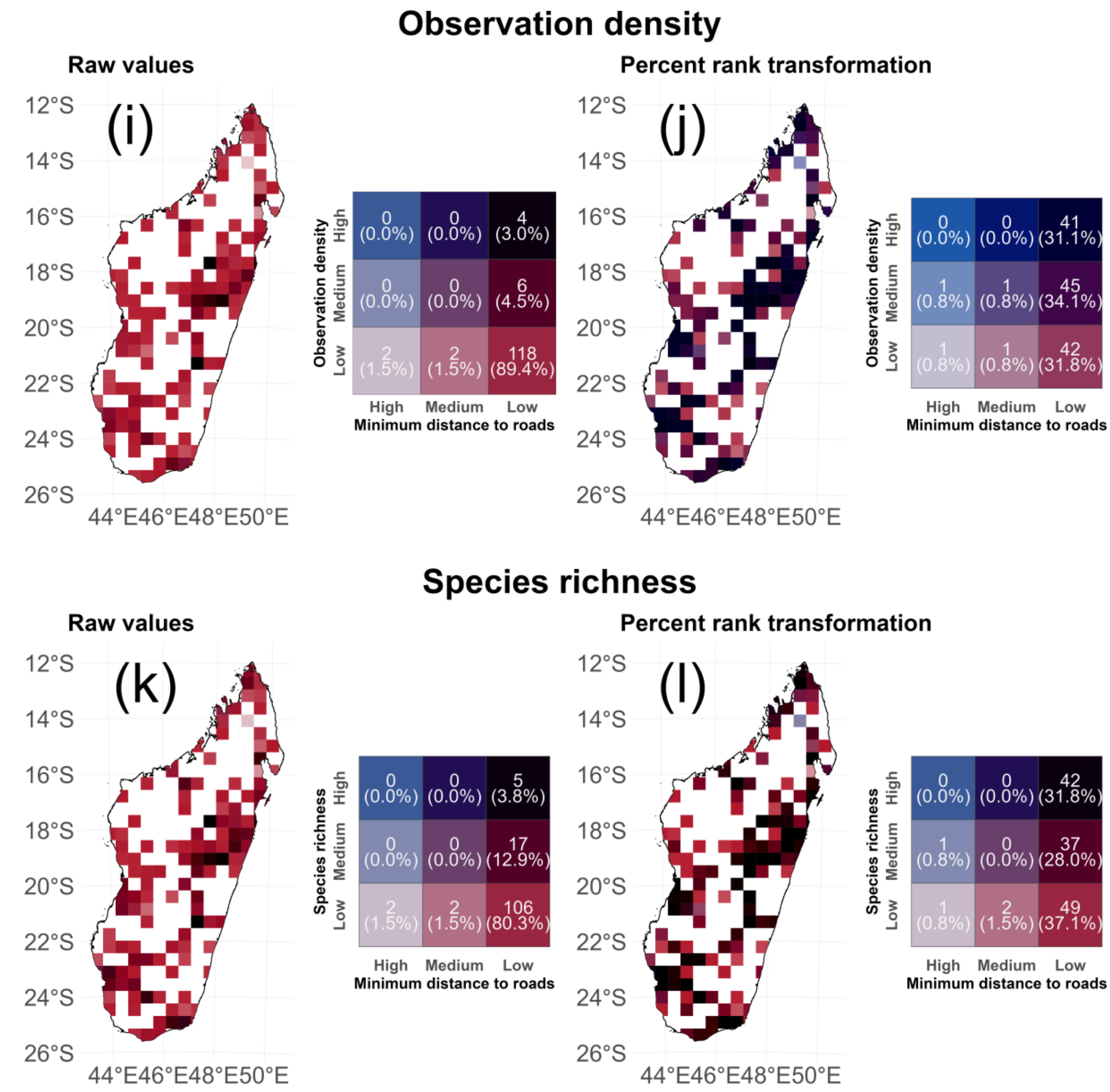


**Supplementary Figure 4. Observation density and species richness at a 10 × 10 km, 25 × 25 km and 50 × 50 km resolutions.** Showing raw values of observation density (OD) and species richness (SR) **(a,c,e,g,i,k)** and percent rank transformation **(b,d,f,h,j,l)**.

**Supplementary Table 4. Metrics calculated for global roads, Madagascar roads and Madagascar KBAs.** For each dataset, the total number of records was reported, along with distance-based metrics measured in kilometers (km). These metrics include: the averaged minimal distance to roads; the standard deviation (SD); the median; the inter-quartile range (IQR), representing the middle 50% of the dataset and calculated as the difference between the 75th percentile (Q3) and the 25th percentile (Q1); the minimum (Min.); and the maximum (Max.). Cumulative proportions are reported at ≤1, ≤2.5, ≤5, and ≤10 km, with the remaining proportion at >10 km. Records located inside KBAs were assigned a distance of 0 and are reported separately within the ≤1 km column (in brackets: proportion inside KBAs + proportion within 1 km).
*26.7% inside KBAs and 13% outside but at ≤1km

| **Data** | **No. of records** | **Mean (km)** | **SD (km)** | **Median (km)** | **IQR (km)** | **Min. (km)** | **Max. (km)** | **≤1km (%)** | **≤2.5km (%)** | **≤ 5km (%)** | **≤10km (%)** | **>10km (%)** |
|---|---|---|---|---|---|---|---|---|---|---|---|---|
| Global roads | 9,216,331 | 1.77 | 20.0 | 0.27 | 0.86 | ~0 | 3425 | 76.9 | 88.9 | 94.6 | 97.5 | 2.52 |
| MDG roads | 4,071 | 1.98 | 2.44 | 1.54 | 1.89 | ~0 | 34.9 | 40.4 | 77.1 | 93.0 | 97.7 | 2.31 |
| MDG KBAs | 4,071 | 10.6 | 17.6 | 1.70 | 14.2 | 0 | 78.8 | 39.7* | 53.5 | 64.8 | 71.9 | 28.1 |

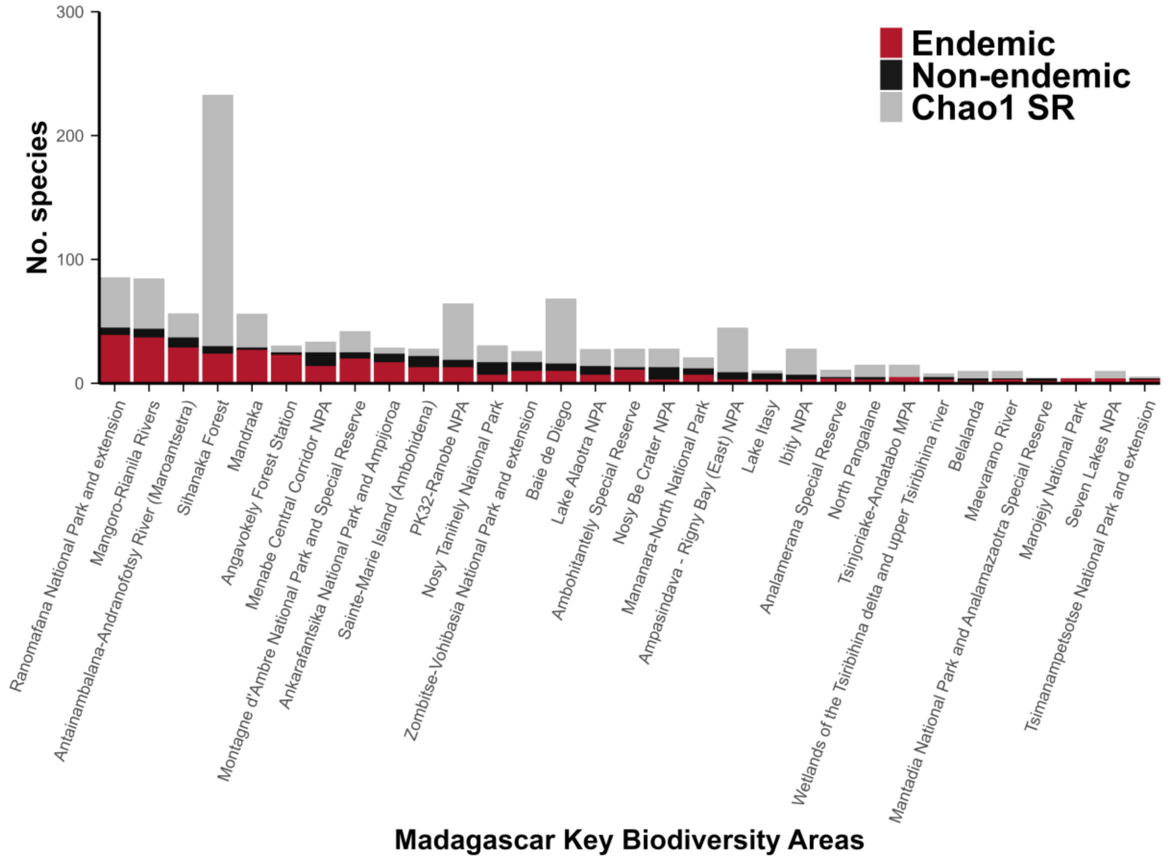


**Supplementary Figure 5. Proportion of Malagasy bee endemics and Chao1 estimated species richness per KBA.**

For visualization purposes, we limited the KBAs shown in the graph to those with more than 3 species records. Most KBAs are highly endemic, with the red bars almost entirely covering the number of observed species. Chao1 estimated species richness (**Chao 1 SR**) is shown for each KBA, highlighting Sihanaka Forest as having particularly high Chao1 values, indicating the presence of rare species recorded within this KBA. The average endemic proportion across all KBAs is 49.5%, although this value is based only on the available records and may increase as additional KBAs are sampled.

### References


Almeida, E. A. B., Bossert, S., Danforth, B. N., Porto, D. S., Freitas, F. V., Davis, C. C., Murray, E. A., Blaimer, B. B., Spasojevic, T., Ströher, P. R., Orr, M. C., Packer, L., Brady, S. G., Kuhlmann, M., Branstetter, M. G., & Pie, M. R. (2023). The evolutionary history of bees in time and space. *Current Biology*, *33*(16), 3409-3422.e6. https://doi.org/10.1016/j.cub.2023.07.005

BirdLife International (2025). The World Database of Key Biodiversity Areas. Developed by the KBA Partnership: BirdLife International, International Union for the Conservation of Nature, Amphibian Survival Alliance, Conservation International, Critical Ecosystem Partnership Fund, Global Environment Facility, Re:wild, NatureServe, Rainforest Trust, Royal Society for the Protection of Birds, Wildlife Conservation Society and World Wildlife Fund. Available at www.keybiodiversityareas.org. [Accessed 17/11/2025].

Bossert, S., Freitas, F. V., Pauly, A., Zhu, G., Crowder, D. W., Orr, M. C., Dorey, J. B., & Murray, E. A. (2025). Phylogeny, antiquity, and niche occupancy of Trinomia (Hymenoptera: Halictidae), an Afrotropical endemic genus of Nomiinae. *Molecular Phylogenetics and Evolution*, *204*, 108273. https://doi.org/10.1016/j.ympev.2024.108273

Danforth, B. N., Eardley, C., Packer, L., Walker, K., Pauly, A., & Randrianambinintsoa, F. J. (2008). Phylogeny of Halictidae with an emphasis on endemic African Halictinae. *Apidologie*, *39*(1), 86–101. https://doi.org/10.1051/apido:2008002

Eardley, C., & Urban, R. (2010). Catalogue of Afrotropical bees (Hymenoptera: Apoidea: Apiformes). *Zootaxa*, *2455*(1). https://doi.org/10.11646/zootaxa.2455.1.1

Freitas, F. V., Branstetter, M. G., Franceschini-Santos, V. H., Dorchin, A., Wright, K. W., López-Uribe, M. M., Griswold, T., Silveira, F. A., & Almeida, E. A. B. (2023). UCE phylogenomics, biogeography, and classification of long-horned bees (Hymenoptera: Apidae: Eucerini), with insights on using specimens with extremely degraded DNA. *Insect Systematics and Diversity*, *7*(4), 3. https://doi.org/10.1093/isd/ixad012

GADM (2025). GADM database of Global Administrative Areas, version 4.1. https://gadm.org

iNaturalist. (2026a). *Parathrincostoma*. Retrieved April 4, 2026, from https://www.inaturalist.org/taxa/606673-Parathrincostoma

iNaturalist. (2026b). *Patellapis*. Retrieved April 4, 2026, from https://www.inaturalist.org/taxa/574257-Patellapis

Meijer, J. R., Huijbregts, M. A. J., Schotten, K. C. G. J., & Schipper, A. M. (2018). Global patterns of current and future road infrastructure. *Environmental Research Letters*, *13*(6), 064006. https://doi.org/10.1088/1748-9326/aabd42

Michener, C. D. (2007). *The bees of the world* (2nd ed). Johns Hopkins university press.

Pauly, A., Brooks, R. W., Nilsson, L. A., Apesenko, Y., Eardley, C. D., Terzo, M., Griswold, T., Schwarz, M., Patiny, S., Munzinger, J., & Barbier, Y. *Hymenoptera Apoidea de Madagascar.* Institute Royal des Sciences Naturelles de Belgique (2001).
